# Transcription factor CCTF1 plays a decisive role in regulating the proteasome activity during archaeal cell division

**DOI:** 10.64898/2026.06.04.729166

**Authors:** Yunfeng Yang, Zixin Geng, Shikuan Liang, Haodun Li, Fan Zhou, Junfeng Liu, Mart Krupovic, Yulong Shen

**Author notes:** These authors contributed equally (Yunfeng Yang, Zixin Geng).

## Abstract

Archaea of the order Sulfolobales exhibit a highly ordered cell cycle progression similar to that of eukaryotes. It has been demonstrated that degradation of the cell division protein CdvB by the proteasome controls the progression of cell division. However, how the proteasome itself is regulated during the cell cycle is not fully understood. Recently, the cyclically expressed ArsR family transcription factor CCTF1 (cell cycle transcription factor 1) was shown to represses expression of PAN (proteasome-activating nucleotidase) in *Sulfolobus acidocaldarius*. Here, using biochemical approaches, comparative transcriptomics and proteomics, we provide further insights into the CCTF1 function and show that it regulates PAN expression in *Saccharolobus islandicus* by binding to an AT-rich palindromic sequence within the *pan* promoter. We also reassessed the role of the cyclically expressed protein kinase aCcrK in regulation of proteasome activity and conclude that CCTF1, rather than aCcrK, is the primary factor controlling the proteasome activity. These findings advance our understanding of the proteosome-mediated cell cycle regulation in archaea.

## Introduction

The cell cycle is a highly conserved and tightly regulated process that ensures accurate replication, segregation, and transmission of genetic material to daughter cells, which is fundamental to the survival and proliferation of all living organisms (1–3). Archaea of the order Sulfolobales are characterized by a eukaryotic-like cell cycle with distinct G1 (approximately 5% of the entire cell cycle), S (approximately 30% of the cell cycle), G2 (approximately 60%), and rapid M and D phases (approximately 5%) (3,4). However, unlike eukaryotes, these archaea lack cyclins and cyclin-dependent kinases (CDKs), the core regulators of eukaryotic cell cycle progression (1,2,5). Therefore, understanding how the cell cycle is regulated in Sulfolobales could provide insight into the origin and evolution of the more complex, cyclin-dependent cell cycle regulation mechanisms (3,6–13). Recent research has suggested that archaeal cell cycle is regulated through multiple mechanisms(14), operating at the levels of transcription, proteosome-mediated protein degradation, and post-translational modification, namely phosphorylation (7,8,10–12,15–17). However, many aspects of archaeal cycle regulation, including regulation of the proteosome activity remain elusive.

Cell division is the final and critical step of the cell cycle, ensuring equal distribution of genetic material and cellular components to daughter cells. The cell division of Sulfolobales relies on the ESCRT-III machinery, which consists of CdvB (also called ESCRT-III), CdvB1 (ESCRT-III-1), and CdvB2 (ESCRT-III-2), and the archaea-specific cell division protein CdvA, The cytokinesis starts with CdvA forming a ring at the mid-cell, which defines the cell division plane (18,19). Subsequently, CdvA ring recruits CdvB through an interaction between the E3B motif of CdvA and the broken wing helix domain of CdvB (19,20). Next, CdvB1/B2 are recruited to the CdvB ring, forming a precisely localized cell division ring together with CdvB (10,21). When the cell is ready to divide, the CdvB polymer is disassembled by the AAA-ATPase Vps4, promoting the constriction of the CdvB1/B2 division ring (6,16). Subsequently, monomeric CdvB is degraded rapidly in *S. acidocaldarious* or gradually in *Sa. islandicus* by the 20S proteasome (6,21–23). Notably, in *S. acidocaldarius*, CdvB degradation is prerequisite for the CdvB1/B2 ring constriction, whereas in *Sa. islandicus*, CdvB disassembly drives the initial membrane constriction, with CdvB1 and CdvB2 playing a key role only at the later stages of cytokinesis (6,16). These differences highlight that even closely related organisms from different genera within the order Sulfolobales can exhibit distinct mechanistic features.

The proteasome, which consists of a proteolytic core particle and a regulatory ATPase, is highly conserved in archaea (24). Members of the order Sulfolobales encode one α subunit and two β subunits, which together form the 20S core particle (25). To activate the proteolytic activity of proteosome, the proteasome-activating nucleotidase (PAN) stimulates the opening of the proteasome α-subunit gate, thereby enabling the degradation of the target proteins (23). The proteasome is essential in *Sa. islandicus*, and none of the genes encoding the proteasomal subunits could be deleted or disrupted (26). Recently, it has been demonstrated that proteasome plays an indispensable role in cell division in Sulfolobales: it mediates the degradation of the key cell division protein CdvB, which is a prerequisite for cytokinesis in both *S. acidocaldarious* and *Sa. islandicus* (6,16). Furthermore, it has been shown that aCcrK (formerly known as ePK2) phosphorylates the α subunit of the 20S proteasome, thereby inhibiting proteasome assembly and regulating proteasome activity (12). Overexpression of aCcrK led to a coherent reduction in cellular proteasome activity and accumulation of cell division proteins. In another study, it was suggested that cell cycle transcription factor 1 (CCTF1) coordinates the cell division in *S. acidocaldarious* by regulating the expression of *pan* (11). However, the precise mechanism by which CCTF1 regulates transcription and whether the proteasome of *Sa. islandicus* is controlled in a similar manner remained unclear.

In this study, we investigate the roles of CCTF1 and aCcrK in modulating the proteasome activity. Through transcriptomic analysis and EMSA experiments, we identified the target genes of CCTF1 as well as the specific binding motifs. By proteomic analyses of CCTF1 or aCcrK overexpressing cells, we have further identified the target proteins regulated by the proteasome. Our results clearly demonstrate that the expression of PAN is specifically regulated by CCTF1 which, in turn, controls the proteosome-medicated degradation of the cell division ring. The results also clarify that aCcrK regulates cell division not by influencing the degradation of the CdvB ring or reduction of proteasome activity but by inhibiting the formation of the CdvB ring. Our study provides important insights into the proteosome-mediated cell cycle regulation mechanism in Sulfolobales.

## Materials and Methods

### Strains, growth conditions and transformation of the strains

*Sa. islandicus* E233S was aerobically grown with shaking at 150 rpm at 76 ℃ in MSTV medium, which contained mineral salts (M), 0.2% (wt/vol) sucrose (S), 0.2% (wt/vol) tryptone (T), and a mixed vitamin solution (200×V). The mutant strain E233S (REY15AΔ*pyrEF*Δ*lacS*) was cultivated in STVU medium with an additional 0.01% (wt/vol) uracil (U). The pH of the culture was adjusted to 3.3 using sulfuric acid, following previously published methods (8,27). MSTV medium was utilized for selecting uracil prototrophic transformants. Culture plates were prepared using gelrite (0.8% [w/v]) by combining 1 × MSTV with an equal volume of 1.6% gelrite. MATV medium containing 0.2% (wt/vol) arabinose (A) was employed for inducing protein overexpression. The plasmids and strains employed in this study are listed in Supplementary Table S1, respectively.

### Bright-field microscopy

For bright-field microscopy analysis, 5 μl of cell suspension at specified time points were observed using a NIKON TI-E inverted fluorescence microscope (Nikon, Japan) in differential interference contrast (DIC) mode.

### Transmission electron microscopy

For negative-staining TEM analysis, cells were stained with 0.3% (w/v) uranyl acetate and observed under an electron microscope (ZEISS crossbeam 550, Germany) operated at 30 kV.

### Cell cycle synchronization

The *Sa. islandicus* E233S strains and *S. acidocaldarius* DSM639 were synchronized following established protocols (8,13,28,29). Initially, the cells were grown aerobically at 75℃ with shaking at 145 rpm in 30 ml of STVU medium. Once the OD_600_ reached 0.6-0.8, the cells were transferred to 100 ml STVU medium with an initial estimated OD_600_ of 0.05 and cultured under the same conditions. Upon reaching an OD_600_ of 0.15-0.2, acetic acid was introduced at a final concentration of 6 mM to arrest the cells at the G2 phase of the cell cycle after 6 hours of treatment. Subsequently, the cells were harvested by centrifugation at 3,000 *g* for 10 minutes at room temperature to eliminate the acetic acid, followed by two washes with 0.7% (w/v) sucrose. The cells were then resuspended in 100 ml of pre-warmed STVU medium and cultured as previously described for further analysis. Flow cytometry was used to analyze the cell cycle, with samples collected at specific time points for subsequent Western blotting or RT-qPCR analysis.

### Flow cytometry analysis

The cell cycle of synchronized E233S cells and strains overexpressing CCTF1 or aCcrK was analyzed using flow cytometry with an ImageStreamX MarkII quantitative imaging analysis flow cytometer from Merk Millipore, Germany. Cell samples were fixed with 70% ethanol for at least 12 hours at each time point and stained with Supergreen. Data was collected for a minimum of 500,000 cells per sample and analyzed using IDEAS data analysis software.

### Protein expression and purification

Recombinant CCTF1 and point mutants with a C-terminal His-tag were expressed in *Escherichia coli* BL21(DE3)-RIL cells. The protein expression was induced during the logarithmic growth phase (OD_600_=0.4∼0.8) by adding 0.5 mM isopropyl-thiogalactopyranoside (IPTG), followed by cultivation at 37 ℃ for 4 hours. The cells were then collected by centrifugation, and the cell pellet was resuspended in buffer A (50 mM Tris-HCl pH 8.0, 300 mM NaCl, 5% Glycerol). After lysing the cells by sonication, the cell extract was clarified by centrifugation at 13,000 × *g* for 20 minutes at 4℃. The supernatant was incubated at 70℃ for 20 minutes, centrifuged again at 12,000 *g* for 20 minutes, and filtered through a 0.45 μm membrane filter. The samples were then loaded onto a Ni-NTA agarose column pre-equilibrated with buffer A and eluted with a linear imidazole gradient (40-300 mM imidazole) in buffer A. The fractions were pooled, concentrated to 1 ml, and further purified by size exclusion chromatography (SEC) using a Superdex 200 column with buffer A. Finally, the samples were dialyzed in a storage buffer containing 50 mM Tris-HCl pH 7.4, 100 mM NaCl, 1mM DTT, 0.1 mM EDTA, and 50% glycerol. The protein concentrations were determined using the Bradford protein assay kit (Beyotime), and the purity was analyzed by 15% SDS-PAGE stained with Coomassie blue.

### Western blotting

The expression levels of CCTF1 in synchronized cells were analyzed using Western blotting. Approximately 2×10_8_ cells were collected at specific time points for each sample and subjected to SDS-PAGE analysis on a 15% gel. The separated proteins were then transferred onto a PVDF membrane. Specific bands were detected using chemiluminescence with Kermey ECL Super Western Blotting Detection Reagents (KermeyBiotech, Zhengzhou, China) as per the manufacturer’s instructions. Primary antibodies against TBP was generated in rabbits by HUABIO (Hangzhou, Zhejiang, China). Antibodies against CCTF1 was produced by AtaGenix Laboratories Co., Ltd. (Wuhan), using purified recombinant proteins(SiRe_1806) from E. coli in rabbits. The primary antibody against the His-tag was purchased from KermeyBiotech (Zhengzhou, China). Goat anti-rabbit antibodies (KermeyBiotech, Zhengzhou, China) conjugated with peroxidase were utilized as secondary antibodies.

### Electrophoretic mobility shift assay (EMSA)

The binding capacity of CCTF1 and its point mutants was assessed using a DNA substrate consisting of a 100 nucleotide (nt) strand labeled with 5’-FAM at the *pan* gene and other gene promoters of *Sa. islandicus* (Supplementary Table S2). The DNA substrates were amplified using specific primers and cloned into pUC19 plasmids at EcoRI and HindIII sites, resulting in a series of pUC19 plasmids with different promoters. FAM-labeled substrates were generated by PCR using FAM-pUC19-EcoRI-F/Specific gene promoter-R primers and various pUC19 plasmids as templates (30). For the binding assay, a 20 μl reaction mixture containing 10 nM dsDNA, 50 mM Tris-HCl (pH 7.4), 5 mM MgCl_2_, 20 mM NaCl, 50 μg/mL BSA, 1 mM DTT, 5% glycerol, and varying concentrations of purified proteins was prepared. The reaction mixture was incubated at 37 ℃ for 30 minutes before being loaded onto a 12% native polyacrylamide gel. Following electrophoresis in 0.5×TBE buffer, the gel was visualized using an Amersham ImageQuant 800 biomolecular imager (Cytiva).

### Transcriptome analysis

Strains of Sis/pSeSD and Sis/pSeSD-CCTF1, Sis/pSeSD-aCcrK were cultured in ATV medium following the described conditions. For transcriptomic analysis, the culture was inoculated with an initial OD_600_ of 0.1. After 12 hours of cultivation, the cells were pelleted at 6,000 × *g* for 10 minutes. The pellet was then resuspended in 1 ml of PBS buffer, pelleted again, and stored at -80℃. Total RNA was extracted using the Trizol reagent (Ambion, Austin, TX, USA) and the RNA Nano 6000 Assay Kit of the Bioanalyzer 2100 system (Agilent Technologies, CA, USA) was used to assess the total amounts and integrity of the RNA. Transcriptomic analysis was carried out by Novogene (Beijing, China), with approximately 3 μg of high-quality RNA per sample used for the construction of RNA-Seq libraries. To begin, mRNA was purified from total RNA using probes to remove rRNA. First strand cDNA was then synthesized using a random hexamer primer and M-MuLV Reverse Transcriptase, followed by RNaseH treatment to degrade the RNA. In the DNA polymerase I system, dUTP was utilized to replace dTTP in dNTP to synthesize the second strand of cDNA.

Exonuclease/polymerase activities were used to convert remaining overhangs into blunt ends, followed by adenylation of the 3’ ends of DNA fragments. Adaptors with hairpin loop structures were ligated to prepare for hybridization. Finally, the USER Enzyme was employed to degrade the second strand of cDNA containing U. To select cDNA fragments of a desired length range of 370∼420 bp, the library fragments were purified using the AMPure XP system from Beckman Coulter in Beverly, USA. Following PCR amplification, the product was further purified with AMPure XP beads to obtain the final library. Subsequently, the libraries were sequenced utilizing the Illumina NovaSeq 6000 platform. The clean reads obtained were then aligned to the reference genome sequence of *Sa. islandicus* REY15A (31). The resulting data was subjected to Fragments Per Kilobase of transcript sequence per Million base pairs sequenced (FPKM) analysis to ascertain the expression levels of all genes within the *Sa. islandicus* genome. Differential expression analysis of the genome (comparing over-expression of aCcr2 or aCcr3 versus an empty vector) was carried out using the DEGSeq R package. To ensure statistical reliability, the resulting P-values were adjusted using the Benjamini and Hochberg’s method for controlling the false discovery rate. A threshold of padj<0.05 and |log2(foldchange)| > 1 was set to determine significantly differential expression.

### Quantitative proteomics

Proteomics analysis was performed at Bioprofile Technology Co., Ltd (Shanghai, China). The specific procedure is as follows:

#### Protein extraction

An appropriate amount of SDT lysis buffer (2% SDS, 100 mM DTT, 100 mM Tris-HCl, pH 7.6) was added to each sample, which was then transferred to an Eppendorf (EP) tube. The samples were incubated in a boiling water bath for 3 min, sonicated for 2 min, and centrifuged at 16,000 *g* for 20 min at 4 °C. The supernatant was collected, and protein quantification was performed using the BCA (bicinchoninic acid) assay. An appropriate amount of protein from each sample was mixed with 5× loading buffer at a ratio of 5:1 (v/v), incubated in a boiling water bath for 5 min, and subjected to 8-16% SDS- PAGE.

#### Protein digestion

An appropriate amount of protein from each sample was digested using the FASP (Filter-Aided Sample Preparation) method (32), as follows: DTT (Dithiothreitol) was added to each sample to a final concentration of 100 mM, followed by incubation in a boiling water bath for 5 min and cooling to room temperature.200 µL of UA buffer (8 M urea, 150 mM Tris- HCl, pH 8.0) was added and mixed, and the mixture was transferred to a 10 kDa ultrafiltration tube and centrifuged at 12,000 *g* for 15 min.200 µL of UA buffer was added, and the tube was centrifuged at 12,000 *g* for 15 min; the filtrate was discarded.100 µL of IAA solution (50 mM IAA in UA buffer) was added, shaken at 600 rpm for 1 min, incubated in the dark at room temperature for 30 min, and centrifuged at 12,000 *g* for 10 min.100 µL of UA buffer was added and centrifuged at 12,000 *g* for 10 min; this step was repeated twice. A amount of 100 µL NH₄HCO₃ buffer was added and centrifuged at 14,000 *g* for 10 min; this step was repeated twice.40 µL of trypsin buffer (6 µg trypsin in 40 µL NH₄HCO₃ buffer) was added, shaken at 600 rpm for 1 min, and incubated at 37 °C for 16-18 h. The collection tube was replaced, and the tube was centrifuged at 12,000 *g* for 10 min to collect the filtrate. The digested peptides were desalted using a C18 cartridge and vacuum- freeze dried. The desalted dried peptides were reconstituted in 0.1% formic acid (FA), and peptide concentration was determined for subsequent LC- MS analysis.

#### DIA (data- independent acquisition) mass spectrometry data acquisition

An appropriate amount of peptides from each sample was separated using a Vanquish Neo UHPLC system (Thermo Scientific). Mobile phase A: 0.1% formic acid in water Mobile phase B: 0.1% formic acid in 80% acetonitrile aqueous solution. The column was equilibrated with 96% mobile phase A. Samples were loaded onto a trap column (PepMap Neo 5 µm C18, 300 µm × 5 mm; Thermo Scientific) and separated on an analytical column (μPAC Neo High Throughput column; Thermo Scientific) using a gradient elution as follows:0-0.1 min: linear increase from 4% to 6% B; 0.1-1.1 min: linear increase from 6% to 12% B;1.1-4.3 min: linear increase from 12% to 22.5% B;4.3-6.1 min: linear increase from 22.5% to 45% B;6.1-8.0 min: held at 99% B. Following separation, DIA analysis (33) was performed on an Orbitrap Astral mass spectrometer (Thermo Scientific) with a total run time of 8 min. Parameters: Electrospray voltage: 2.2 kV, Detection mode: positive ion, Precursor scan range: 380-980 m/z,MS1 resolution: 240,000,MS1 AGC target: 500%,MS1 maximum IT: 3 ms,MS2 resolution: 80,000,MS2 AGC target: 500%,MS2 maximum IT: 3 ms, RF lens: 40%,MS2 activation type: HCD, Isolation window: 2 Th, Normalized collision energy: 25%,Cycle time: 0.6 s.

#### DIA mass spectrometry data analysis

All mass spectrometry data were processed and integrated using DIA- NN software for database searching and DIA- based protein quantification(33,34). The database used was: uniprotkb- *Saccharolobus islandicus* (species) [43080] - 27721- 20260128; fasta- https://www.uniprot.org/taxonomy/43080

#### Data analysis and bioinformatics analysis

To ensure the validity and accuracy of subsequent bioinformatics and statistical analyses, we first filtered the experimental data in the protein identification table following common criteria. Proteins were retained only if they had non-missing values in at least 50% of the samples within their corresponding group. The remaining missing values were then subjected to data imputation before statistical analysis. Proteins meeting the criteria of fold change > 1.5 (either up- or down-regulation) and P-value < 0.05 were defined as significantly differentially expressed proteins.

### Quantitative reverse transcription PCR (RT-qPCR)

Quantitative reverse transcription PCR (RT-qPCR) was utilized to analyze changes in the expression levels of aCcr1 and its homologous genes. Samples were collected from E233S strains during the logarithmic growth phase and after synchronization at the corresponding time points. Total RNA extraction was carried out using SparkZol (SparkJade Co., Shandong, China). First-strand cDNAs were synthesized from the total RNA following the protocol of the First Strand cDNA Synthesis Kit (Accurate Biotechnology Co., Hunan, China) for RT-qPCR analysis. The resulting cDNA samples were used to measure the mRNA levels of the target genes through qPCR using the SYBR Green Premix Pro Taq HS qPCR Kit (Accurate Biotechnology Co., Hunan, China) and gene-specific primers (Supplementary Table S2). PCR was conducted in a CFX96TM (Bio-Rad) instrument with the following steps: denaturation at 95℃ for 30 seconds, followed by 40 cycles of 95℃ for 5 seconds and 60℃ for 30 seconds. The relative quantities of mRNAs were determined using the comparative Ct method with the 16S rRNA gene as the internal reference. All qPCR primers were verified to have an amplification efficiency within the range of 90% to 110%.

### Fluorescence microscopy

Fluorescence microscopy analysis was performed as previously described (16,27,28,35). Briefly, cells were collected and pelleted down at 3,000 *g* for 5 min, and resuspended in 300 μl PBS buffer (137 mM NaCl, 2.7 mM KCl, 10 mM Na_2_HPO_4_ × 12H_2_O, 2 mM KH_2_PO_4_). The cells were fixed by adding 700 μl of cold absolute ethanol at 4℃ for 2 h. Then the cells were pelleted down and washed with PBST buffer (PBS plus 0.05% Tween-20) for 3 times at 3,000 *g* for 3 min. The primary antibody against CdvB by HUABIO (Hangzhou, Zhejiang, China) was added (dilution of 1:1000 in PBST buffer) and incubated at 4℃ overnight. The cells were then pelleted down and washed with PBST buffer for 3 times at 3,000 *g* for 5 min. Goat anti-rabbit IGG Alexa Fluor® 568, Invitrogen™ (Thermo Fisher Scientific, USA) was added (dilution of 1:1000 in PBST) and incubated at room temperature for 2 h. Then the cells were pelleted down and washed with PBST buffer for 3 times at 3,000 *g* for 5 min. The cells were finally resuspended in PBS buffer containing DAPI (4’, 6-diamidino-2-phenylindole) to stain the DNA. After 30 min of staining, the samples were observed under Zeiss LSM 880 confocal microscope (Germany).

## Results

### Cyclically expressed gene *cctf1* is indispensable for cell viability

Previous studies indicated that proteasome is involved in the degradation of cell division protein CdvB (6). More recently, it was reported that the proteasome regulatory subunit PAN (proteasome-activating nucleotidase) is expressed in a cyclic manner at both transcriptional and translational levels, indicating that proteasome activity is regulated cyclically (36). Overexpression of an ArsR (arsenical resistance operon repressor) type transcription factor CCTF1 repressed the expression of *pan* and led to accumulation of CdvB in *S. acidocaldarius*, suggesting that CCTF1 regulates cell division and hence the cell cycle by modulating the cellular proteosome activity (11,36). To elucidate the mechanistic details of CCTF1 cell cycle regulation, we first reevaluated the previously reported transcriptome data of synchronized *Sa. islandicus* population. We found that the transcription of the gene coding for the CCTF1 homolog in *Sa. islandicus*, *sire_1806,* hereinafter denoted *siscctf1*, exhibits an apparent cyclic expression pattern, with peak expression during cytokinesis (Figure S1B)(8). The RT-qPCR analysis confirmed the cyclic expression pattern (Figure 1A and 1B). Notably, the peak expression time point of *siscctf1* was close to those of *cdvB* and *pan*, and ∼0.5 hr later than those of *cdvA* and *aCcrK*. To examine if CCTF1 is expressed cyclically at the protein level, we prepared rabbit antibodies against *Sis*CCTF1, using purified recombinant protein from *E. coli*. The antibody exhibited equivalent efficacy against recombinant CCTF1 proteins from both *Sa. islandicus* and *S. acidocaldarius* (Figure S1C). For unknown reason, the signal could not be detected in the cell extract of *Sa. islandicus* in Western blotting analysis using this antibody. However, a band of expected size was detected in the cell extract of *S. acidocaldarius*. Therefore, we used *S. acidocaldarius* to examine the protein expression during the cell cycle with the anti-CCTF1 antibody. As shown in Figure S1D and S1E, the levels of *Sa*CCTF1 exhibited a cyclic pattern, with a peak at approximately 2.5 hours, coinciding with the cell division phase. The expression levels of *pan* and *cdvB* peaked roughly at same time (Figure S1B). For technical reasons, it remains challenging to distinguish the temporal order of expression for CCTF1, PAN, and CdvB using the current synchronization methodology. In addition, we attempted to knockout *cctf1* using the endogenous CRISPR-Cas based genome editing technology. Despite numerous efforts, we were unsuccessful in generating a knockout strain, suggesting that *cctf1* is essential for cell viability. This result is in agreement with a previous report on the essentiality of *Sa. islandicus* genes (26).

**Figure 1.**
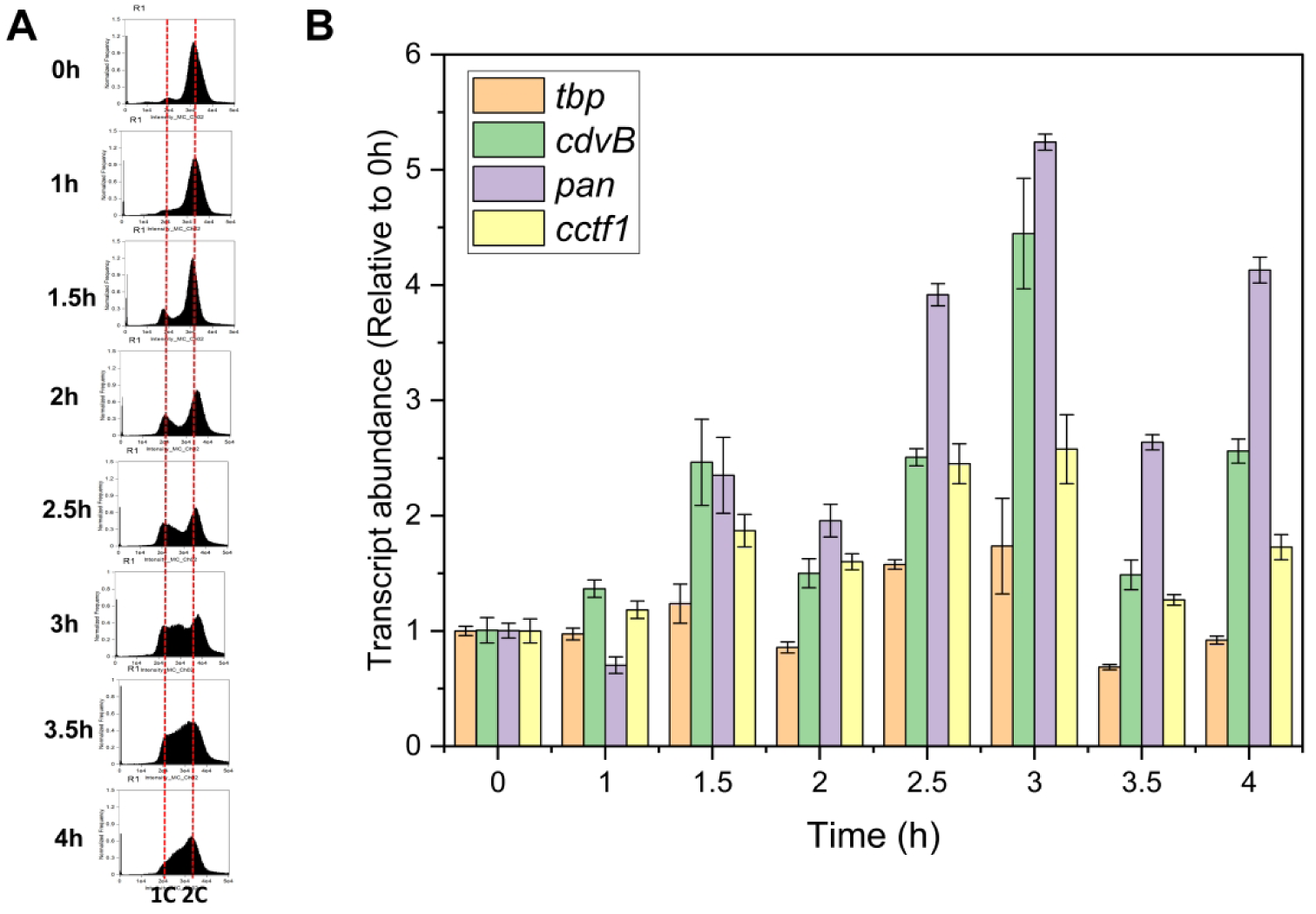
The cell cycle transcription factor CCTF1 displays a cyclic pattern at both the transcriptional and protein levels. (**A**) Flow cytometry profiles of samples of a synchronized *S. islandicus* REY15A (E233S) culture.. The cells were synchronized at G2 phase after treated with 6 mM acetic acid for 6 h before released by removing the acetic acid. The cultures were collected at different time points (0, 1.0, 1.5, 2.0, 2.5, 3.0, 3.5 and 4.0h) and subjected to flow cytometry analysis. ‘1C’ represents one chromosome, ‘2C’ represents two chromosomes. (**B**) RT-qPCR analysis of the transcription of *cdvB*, *cctf1* and *pan* at different cell cycle phases, The 16S rDNA was used as an internal reference and the abundance of *tbp* was set as 1. The experiments were performed in three biological repeats.

### Overexpression of *Saci* CCTF1 leads to severe growth inhibition, abnormal cellular DNA content, and cell death in *Sa. islandicus*

Unexpectedly, our attempts to construct a *Sis*CCTF1 (SiRe_1806) overexpression strain in *Sa. islandicus* were unsuccessful, suggesting that without tight expression regulation (not provided by our expression vector) this protein exhibits cytotoxicity. However, we obtained a *Sa. islandicus* E233S strain overexpressing *Saci*CCTF1. *Saci*CCTF1 and *Sis*CCTF1 share ∼60% amino acid sequence identity (Figure S1A). *Saci*CCTF1 overexpression induced with arabinose severely inhibited cell growth (Figure 2A). The cells exhibited an abnormal flow cytometry profile: after 24 hours of induction, many cells contained more than 2 or less than 1 chromosome copies, the latter corresponding to dead cells (Figure 2D). Consistently, cells displayed abnormal morphology, with many cells being enlarged and occasionally interconnected (Figure 2BC). These results indicate that CCTF1 overexpression affects not only cell division but also chromosome segregation and/or genome replication. In addition, we obtained a strain overexpressing *Saci*CCTF1 with the C-terminal His-tag. This strain exhibited the same phenotype as the strain overexpressing the wild type protein (Figure S1F and S1G), suggesting that addition of the His-tag did not affect the activity of the protein *in vivo.* For this reason, we purified the C-terminally His-tagged wildtype and mutant CCTF1 proteins for electrophoretic mobility shift assay (EMSA; see below).

**Figure 2.**
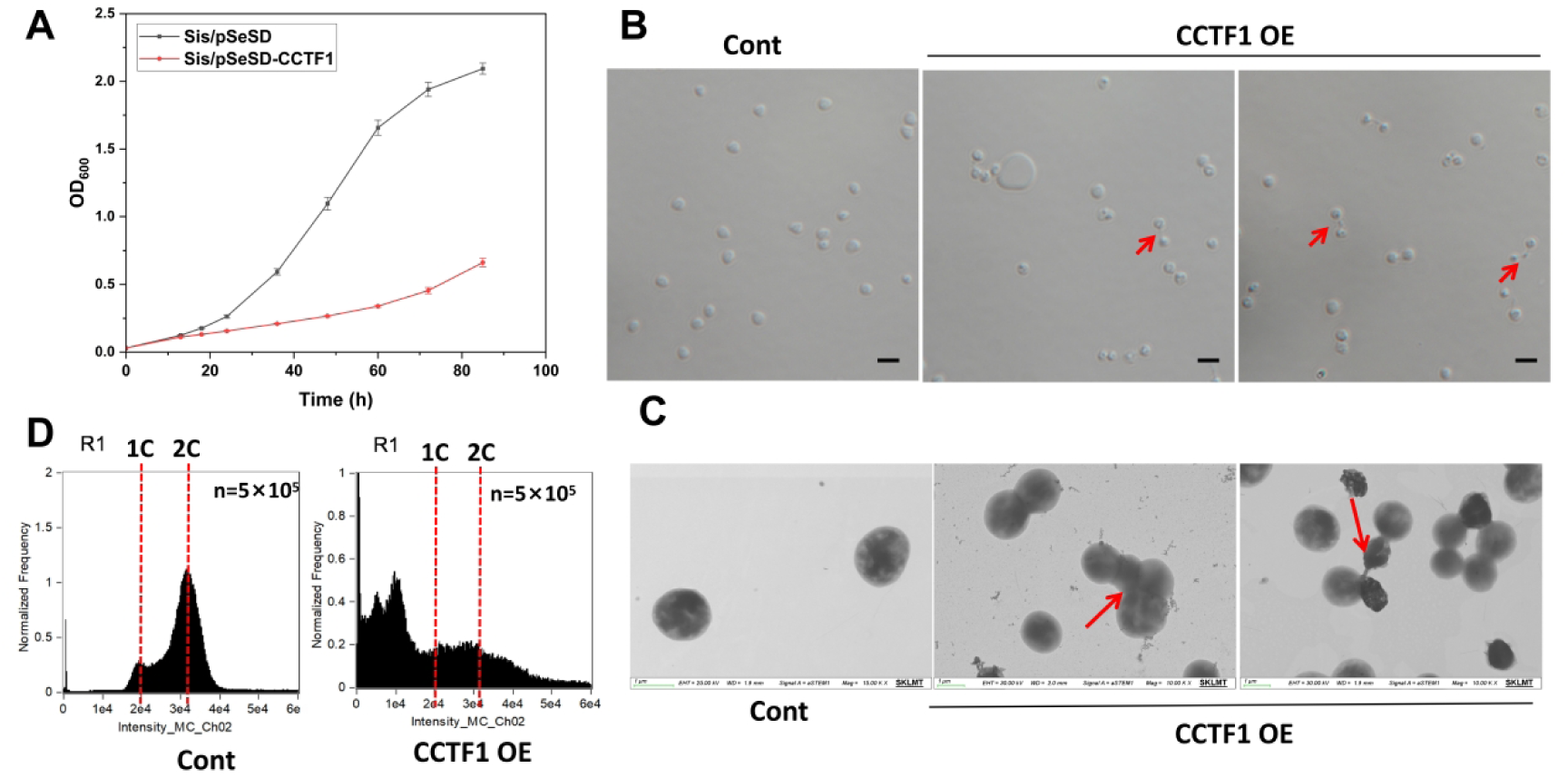
CCTF1 overexpression leads to severe growth inhibition, defects in cell division, and chromosome content abnormality. (**A**) Growth curves of cells over-expressing *Saci*CCTF1. The cells were inoculated into 30 ml induction medium ATV to a final estimated OD_600_ of 0.03. The growth was monitored at OD_600_. Each value was based on data from three independent measurements. Cells harboring the empty plasmid pSeSD were used as a control. (**B**) and (**C**) Morphology of cells overexpressing CCTF1 observed by bright-field microscopy (DIC) (**B**) and transmission electron microscopy (TEM) (**C**). Scale bars: 2μm. The red arrows indicate cell under abnormal division or dead. (**D**) Flow cytometry of cells over-expressing CCTF1. Cells cultured in the induction MTAV medium were taken at 24h for the microscopical and flow cytometry analyses.1C represents one chromosome, 2C represents two chromosomes.

### The transcription of the proteasome regulatory subunit PAN is specifically regulated by CCTF1

To identify genes transcriptionally regulated by CCTF1 and to assess the impact of CCTF1 overexpression on cell division, we conducted a comparative transcriptomic analysis between a *Saci*CCTF1-overexpressing strain and a control strain harboring the empty vector pSeSD (Figure 3A). Samples were collected 12 hours post-arabinose induction and subjected to transcriptome sequencing. A total of 95 and 103 genes were up- and down-regulated more than twofold, respectively (Figure 4A Supplementary Table S3). The proteasome regulatory subunit gene *pan* (*sire_1726*) ranked at the top of the most significantly down-regulated genes. There was no significant change in the expression of genes associated with cell division, with only *cdvB1* showing a slight upregulation (log2FoldChange=1.02; padj=0.005). We therefore speculate that the effect of CCTF1 overexpression on cell division is largely due to severe repression of *pan*.

**Figure 3.**
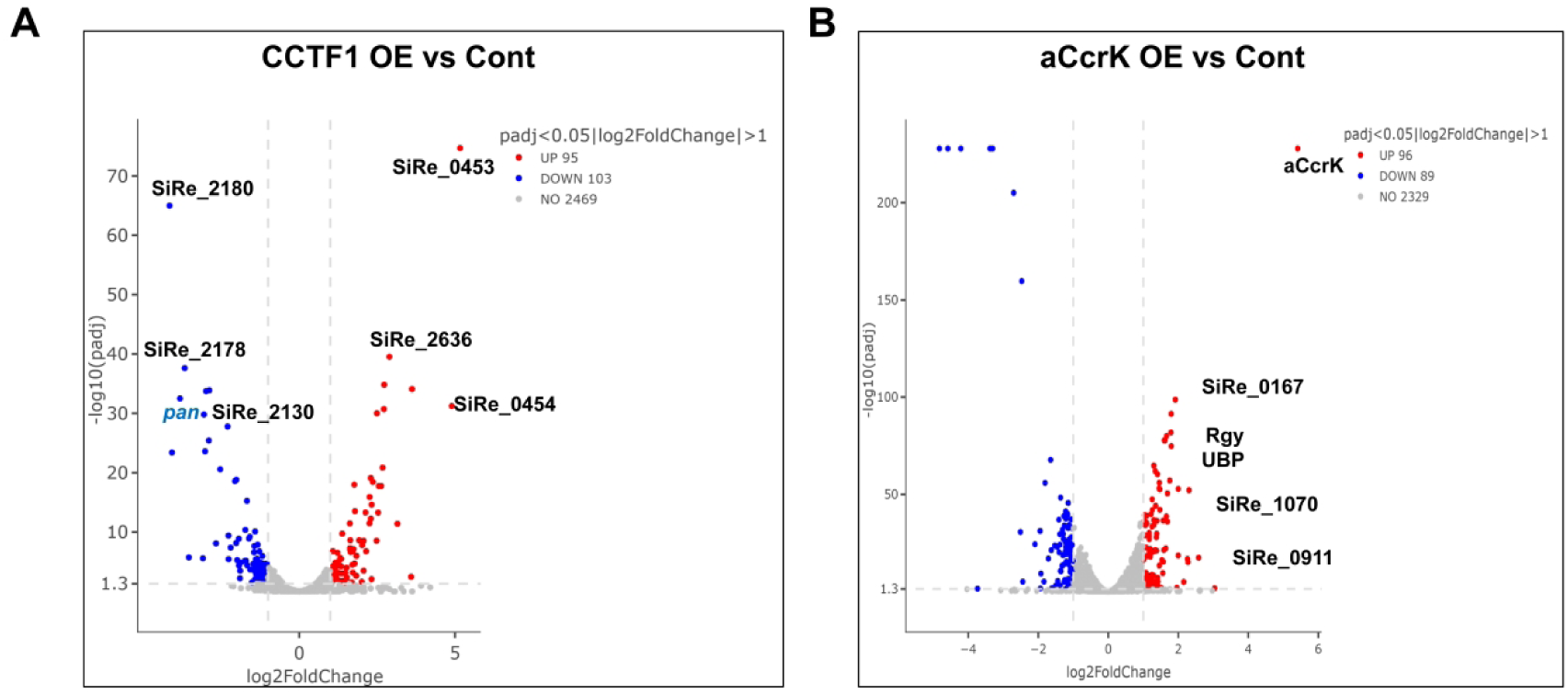
Comparative transcriptomic analysis of CCTF1 and aCcrK overexpression strains. (**A**) and (**B**) Volcano plots of differentially expressed genes in strains Sis/pSeSD-CCTF1 and Sis/pSeSD-aCcrK compared to the control Sis/pSeSD. X-axis, log2fold change in gene expression. Y-axis, significance of fold change. Genes exhibiting > 2-fold (i.e. –1 > log2 > +1) significant upregulation and downregulation are highlighted in red and blue, respectively, whereas those that showed a <2-fold change in differential gene expression or with no significance are shown in gray. The *pan* gene is marked in blue.

**Figure 4.**
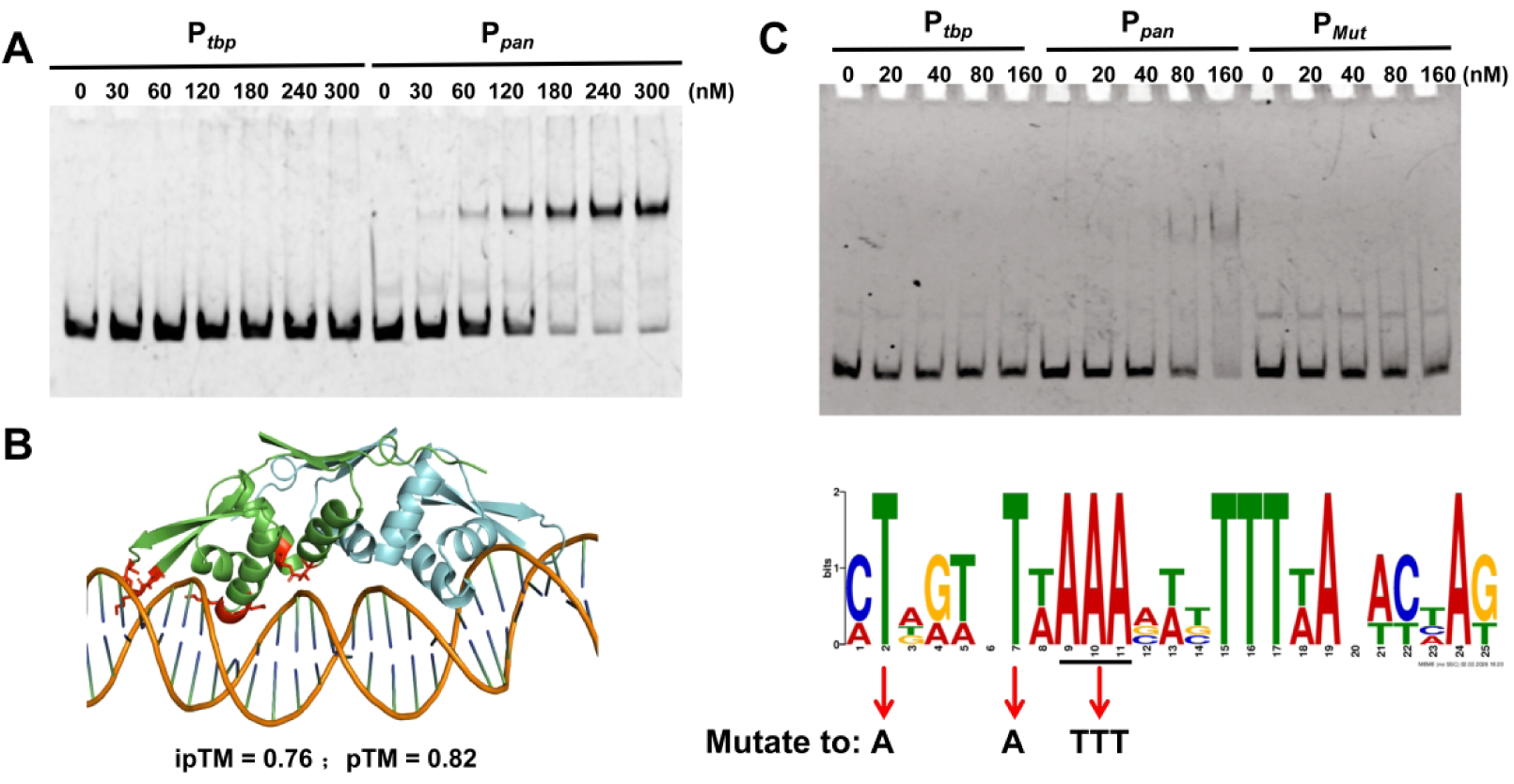
CCTF1 specifically regulates the expression of the proteasome regulatory subunit *pan*. (**A**) EMSA for the DNA binding activity of the CCTF1. P*_tbp_* and P*_pan_* was used as the substrates. Each reaction contained 10 nM of the 5’-FAM labelled probe. (**B**) Model of *Sis*CCTF1 binding to the promoter of *pan* (TGTAAATACTTGTATTAAACATTTTTAGTCTATGTCAAACAAGTATATAT). The structure was predicted using AlphaFold3 (https://alphafoldserver.com/). The ipTM and pTM values (which reflect the significance of the model) are displayed below the corresponding model. Key amino acid binding sites are highlighted in red. (**C**) EMSA for the DNA binding activity of *Sis*CCTF1. P*_tbp_*, P*_pan_* and P*_mut_* were used as the substrates. Each reaction contained 10 nM of the 5’-FAM labelled probe. Potential CCTF1 binding motifs predicted from CCTF1-repressed genes are shown at the bottom. The mutated bases in the P*_mut_* promoter are shown in the figure.

To test whether CCTF1 binds to the promoter of *pan*, we performed an EMSA analysis using a DNA substrate containing the sequence upstream of the translation start site and the 5’ untranslated region (5’ UTR) of *pan*. Purified *Sis*CCTF1 (0, 30, 60, 120, 180, 240 and 300 nM) was incubated with fluorescein (FAM)-labelled DNA substrate (Figure 3A). We found that CCTF1 was able to bind P*_pan_* and the signal intensity increased with the increasing concentration of the protein (Figure 4A). No electrophoretic mobility shift was observed when P*_tbp_* (negative control) was used as the substrate. We also performed EMSA using substrates containing promoter regions of several other downregulated genes coding for DUF5751 domain-containing protein (SiRe_0150), inorganic pyrophosphatase (SiRe_0183), biotin carboxylase (SiRe_0254), oligosaccharyltransferase membrane subunit (SiRe_1041), Alba (SiRe_1125), signal peptidase I (SiRe_1168), and periplasmic component of the ABC-type sugar transport system (SiRe_2180). The results showed that, in addition to *pan*, CCTF1 also specifically bound to P*_sire_0183_*, P*_sire_0254_*, P*_sire_1041_*, P*_sire_1125_* (Alba), and P*_sire_2180_* (Figure S3A). Analysis of the putative promoter sequences bound by CCTF1 allowed us to identify an AT-rich palindromic sequence common to P*_pan_,* P*_sire_0183_*, P*_sire_0254_*, P*_sire_1041_*, P*_sire_2180_*, and P*_sire_1125_* (Figure 4C and S3B and S2C), which is likely the specific binding motif for CCTF1. To test this possibility, we mutated the putative motif by replacing the adenine (A) with thymine (T) and, conversely, mutated T to A in the 5’ half of the sequence, thereby disrupting the palindromic structure. As expected, compared to the substrate containing the wild-type motif, the binding affinity of *Sis*CCTF1 for the mutated sequence was greatly reduced, supporting the hypothesis that the palindromic sites are important for CCTF1 binding (Figure 4C).

CCTF1 belongs to the ArsR family of transcription factors, which adopt the winged helix-turn-helix (wHTH) fold. We used AlphaFold3 to predict the CCTF1 structure with the cognate DNA (Figure 4B). Based on the obtained structural model, we constructed stains carrying the putative DNA-binding site mutants and analyzed their phenotypes. We also purified the recombinant mutant proteins and performed EMSA to verify the importance of the predicted residues for the DNA binding activity of CCTF1. The results suggest that K73, G74, and K76 located within the wing region are crucial for the DNA-binding (Figure S2). When all three of these residues were mutated, the DNA binding was completely abolished (Figure S2B). Consistently, cells overexpressing the mutant proteins displayed normal growth, similar to that of cells carrying the empty plasmid (Figure S2A). The results suggest that DNA binding of CCTF1 is important for its activity *in vivo*.

### CCTF1, rather than aCcrK, plays a pivotal role in regulating proteasome activity

The results presented above support the conclusion that CCTF1 regulates the proteosome activity by transcriptional regulation of the regulatory subunit of the 26S proteosome, thereby controlling cell division. We have recently reported that overexpression of the kinase aCcrK (formerly known as ePK2) in *Sa. islandicus* affected cell division through phosphorylation of the α-subunit of the proteasome, which inhibited the proteasome activity (12). To clarify the roles of CCTF1 and aCcrK in cell division regulation, we analyzed the phenotypes of CCTF1- and aCcrK-overexpressing strains. Flow cytometry profiles of synchronized cells overexpressing CCTF1 and aCcrK showed that cell division was inhibited in both stains (Figure 5). However, enlarged cells with more than two copies of the chromosome and dead cells with fewer than one copy of the chromosome were generated only in the CCTF1 overexpression strain, suggesting abnormal chromosome segregation. In contrast, only large cells with more than two copies of the genome were produced in the aCcrK overexpression strain (Figure 5 and S4B). To further illustrate the functional difference between CCTF1 and aCcrK in proteosome-mediated protein degradation, we determined the growth curves of the aCcrK-overexpressing cells, the CCTF1-overexpressing cells, and wild-type cells treated with Bortezomib, the proteasome inhibitor. The results showed that the growth inhibition caused by aCcrK overexpression is far less pronounced than that caused by CCTF1 overexpression or treatment with the proteasome inhibitor (Figure S4A). These results suggest that the mechanisms underlying the cell division abnormalities induced by the overexpression of these two proteins are different.

**Figure 5.**
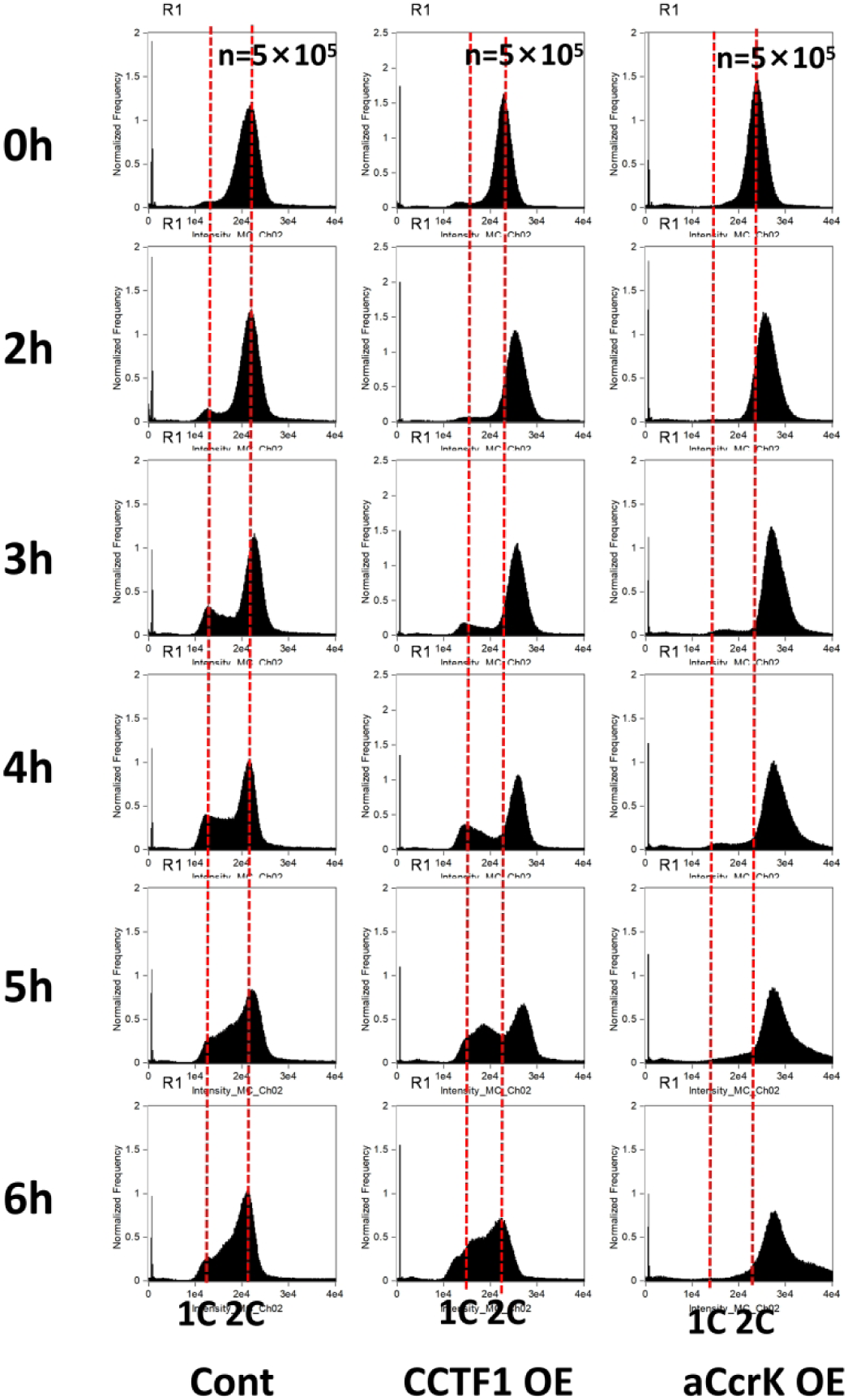
Overexpression of CCTF1 and aCcrK affects cell division in different ways. Flow cytometry profiles of DNA content distribution of cells 0-6 hrs after acetic acid removal. E233S containing the empty plasmid (Sis/pSeSD) was used as a control. 1C represents one chromosome, 2C represents two chromosomes.

To elucidate the specific mechanisms by which CCTF1 and aCcrK overexpression affect cell division, we conducted a proteomic analysis and coupled the obtained results with the comparative transcriptomic data (Figure 3AB, 6AB, and Supplementary Table S3,S4). If expression of either protein affects the proteasome activity (by either inhibiting the expression of the PAN subunit or through phosphorylation of the proteasome α subunit), the changes in the proteome of both strains are expected to be similar to those in the strain treated with the proteasome inhibitor. However, the proteome of the CCTF1 overexpressing cells was similar to that observed following treatment with Bortezomib, consistent with the previous report (6). In particular, the cell division proteins CdvB, CdvB1, and CdvB2 displayed accumulation, suggeststing that PAN is one of the main targets of CCTF1 and CTTF1 is involved in the regulation of proteosome-mediated degradation of the cell division proteins (Figure 6A and 6F). The transcriptomic data of the CCTF1 overexpression cells showed that only CdvB1 was slightly upregulated (log2FoldChange=1.02; padj=0.005), while CdvB and CdvB2 showed no significant changes (Figure 6F), strongly suggesting that their accumulation is indeed caused by the inhibition of the proteasome activity. In contrast, the comparative proteomic analysis of the aCcrK overexpressing cells showed that only CdvB1 displayed higher levels (log2FoldChange=1.82; p value=0.0006), likely due to transcriptional upregulation of *cdvB1* (log2FoldChange=1.30; padj=1.82E-37) (Figure 6F).

**Figure 6.**
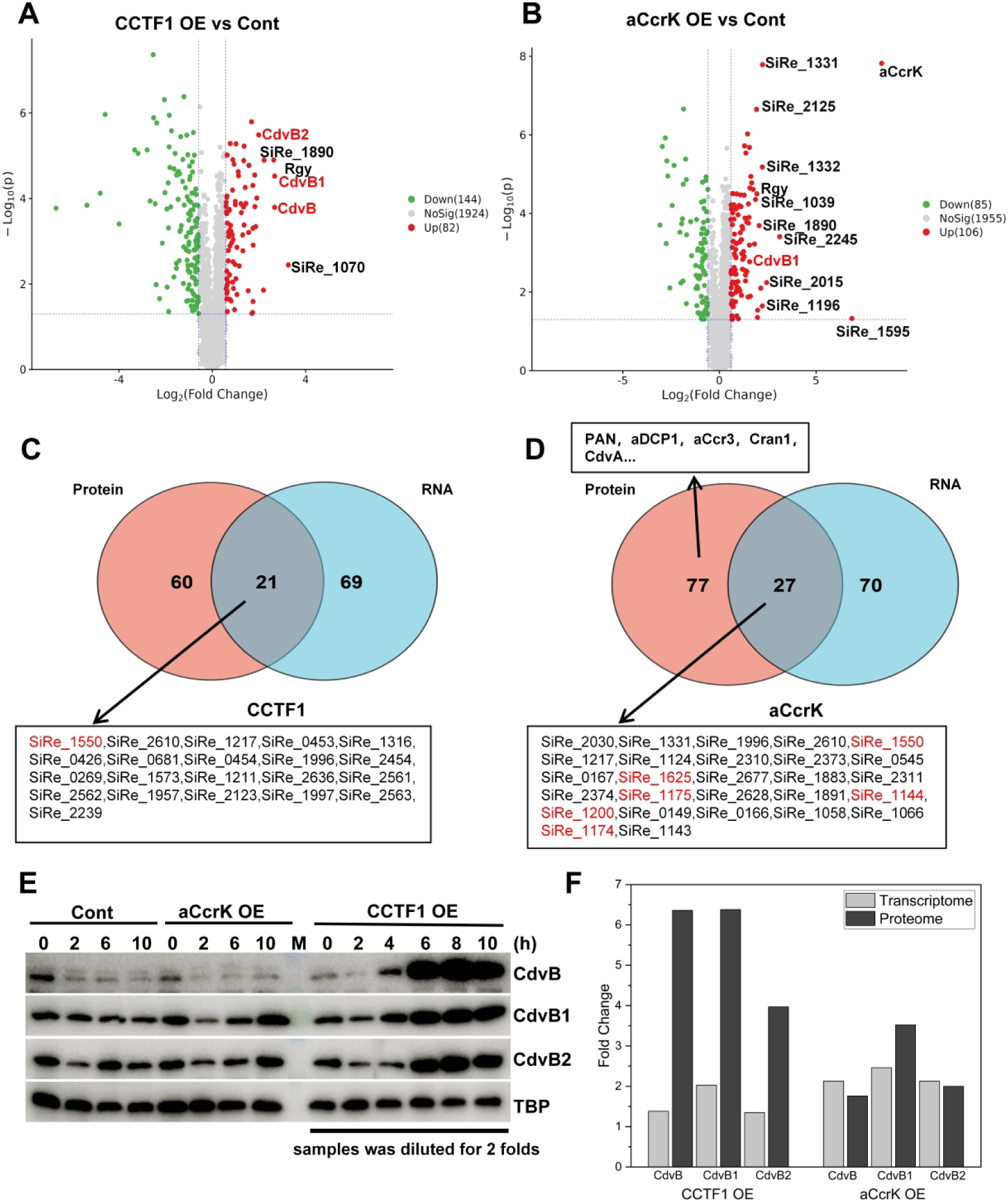
Proteomic Analysis of CCTF1 and aCcrK Overexpression Strains. (**A**) and (**B**) Volcano plots of differentially expressed proteins in strains Sis/pSeSD-CCTF1 and Sis/pSeSD-aCcrK compared to the control Sis/pSeSD. X-axis, log2fold change in protein expression. Y-axis, significance of fold change. Proteins exhibiting > 1.5-fold (i.e. –0.59 > log2 > +0.59) significant upregulation and downregulation are highlighted in red and green, respectively, whereas those that showed a <1.5-fold change in differential protein expression or with no significance are shown in gray.The cell division proteins is marked in red. (**C**) and (**D**)Venn diagram analysis of the genes that are upregulated in both the transcriptome and proteome in the CCTF1 (C) and aCcrK (D) overexpression strains, respectively.Genes marked in red are known to accumulate significantly upon treatment with a proteasome inhibitor(6). (**E**) Verify the expression levels of the cell division proteins by western blotting in CCTF1 and aCcrK overexpression strains. As the high levels of CdvB in the CCTF-overexpressing strain were interfering with signal visualisation in other strains, we diluted the CCTF sample twofold. (**F**) Changes in the expression of cell division proteins in the transcriptomes and proteomes of CCTF1 and aCcrK overexpression strains.

Comparison of the genes and proteins that were up-regulated in the transcriptomic (fold change > 2) and proteomic analyses (fold change > 1.5) in the CCTF1 and aCcrK overexpression strains (Figure 6C and 6D), respectively, showed that the upregulation of proteasome-regulated target proteins in the CCTF1 overexpression strain was not caused by their transcriptional upregulation. By contrast, in the aCcrK overexpression strain, target proteins (such as CdvB, CdvB1, CdvB2, XerA, TFB2, Vps4, etc.) were all upregulated at the transcriptional level (Figure 6C and 6DD). At the same time, the Western blotting experiments confirmed that there is no apparent difference in the level of CdvB, CdvB1 or CdvB2 between the aCcrK-overexpressing cells and the control cells. However, the levels of all three cell division proteins were significantly higher in the CCTF1-overexpressing strain, compared to the control. In particular, the amount of CdvB in CCTF1 overexpressing cells was 50–100 times higher than that in the control strain (Figure 6E). In addition, we compared changes in CdvB protein levels in CCTF1-overexpressing and aCcrK-overexpressing strains with those observed following treatment with the proteasome inhibitor bortezomib. Western blotting results clearly demonstrated that aCcrK overexpression does not lead to CdvB accumulation, whereas CCTF1 overexpression, similar to treatment with the proteasome inhibitor, results in CdvB accumulation (Figure S4C). Meanwhile, we have noted that no significant change in the transcriptional levels of *pan*(log2FoldChange=0.25; padj=0.01997) in the transcriptome of aCcrK-overexpressing cells; however, proteomic data indicated higher levels of PAN(log2FoldChange=1.49; pvalue=0.00018) (Figure 6D). Therefore, proteasome activity would increase, and this was also evident from the Western blotting results on CdvB. In contrast, the levels of CdvB in the aCcrK-overexpressing strain were lower at 4 h and 8 h post-arabinose induction compared to 2 h (Figure S4C).We therefore hypothesize that the cell division deficiency induced by overexpression of aCcrK is unlikely to be due to reduced activity caused by phosphorylation of the proteasome α subunit, but rather due to as of yet unknown mechanism. Furthermore, the results support the previous conclusion that CCTF1 is a transcription factor that regulates the expression of the proteasome regulatory subunit that is crucial for controlling the proteasome activity.

### Overexpression of CCTF1 suppresses CdvB ring disassembly, whereas overexpression of aCcrK impedes CdvB ring formation

To further investigate the mechanisms by which CCTF1 and aCcrK regulate cell division, we performed immunofluorescence localization of CdvB in synchronized cultures undergoing the division phase (i.e., 3 hours after removal of acetic acid). As shown in Figure 5, in the control cells, 7.8% of cells had CdvB rings, whereas in the CCTF1-overexpressing strain, approximately 44.03% of cells had such rings. In contrast, only approximately 0.67% cells had the CdvB rings in the aCcrK overexpressing cells (Figure 7A and 7B). This result suggests that CCTF1 overexpression indeed disturbs CdvB ring disassembly, whereas aCcrK overexpression impedes CdvB ring formation. Notably, in the asynchronous population (20 hours after induction with arabinose), very high proportion (61.07%) of cells with CdvB rings were observed in the CCTF1-overexpressing strain, whereas only 1.48% of the cells displayed CdvB rings in the aCcrK overexpression cells (Figure S5).

**Figure 7.**
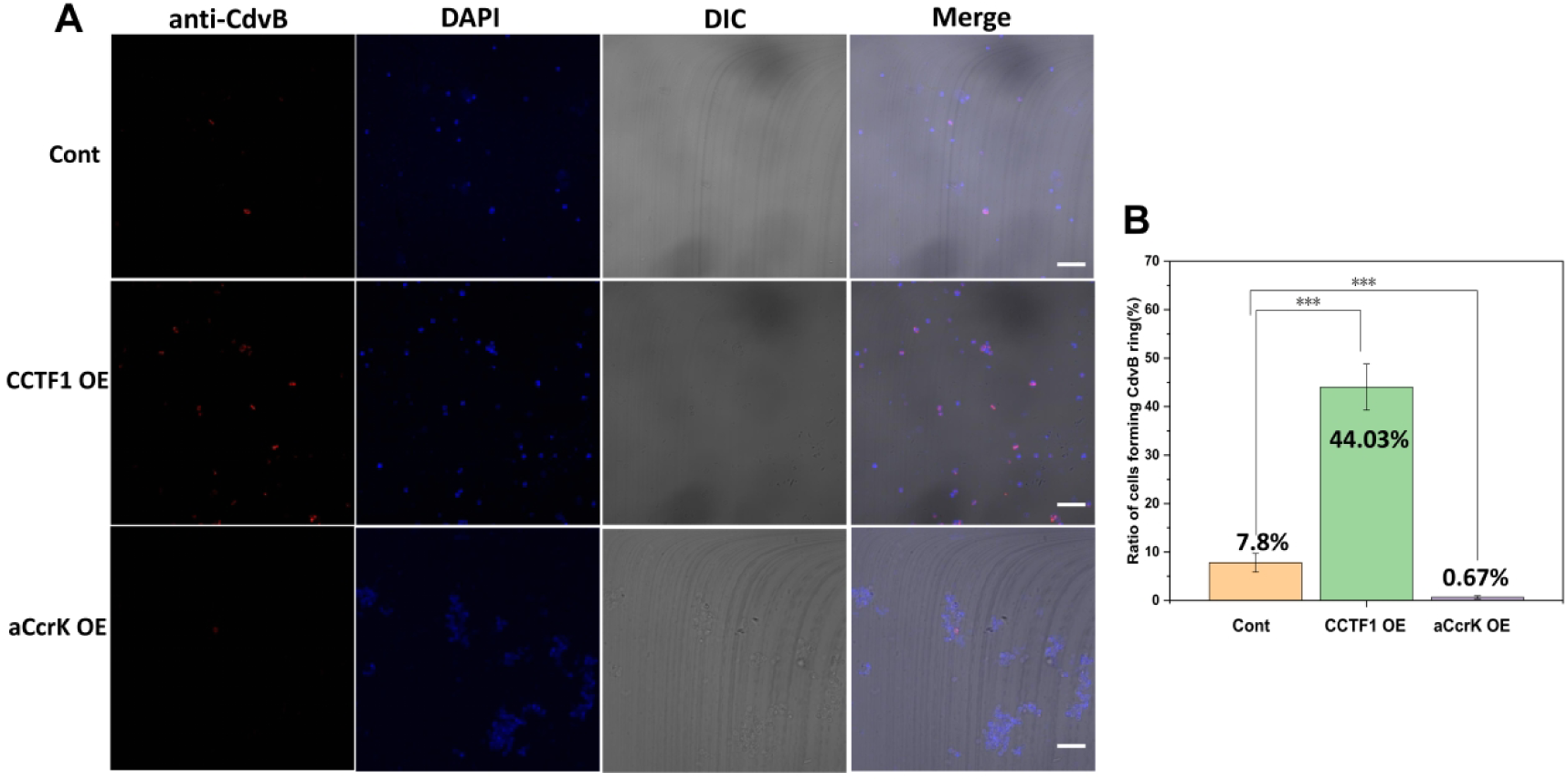
Overexpression of CCTF1 suppressed the degradation of the CdvB ring, whilst overexpression of aCcrK prevented the formation of the CdvB ring. (**A**) Immuno-fluorescence microscopy showing the formation of contractile rings using the primary antibody against ESCRT-III (CdvB) and goat anti-rabbit secondary antibody Alexa Fluor® 568. The DNA was stained with DAPI. Shown are representative images. This samples were obtained from cells 3 hours after synchronisation, as shown in Figure 6. (**B**) Statistical analysis of the proportion of CdvB ring formation in the control strain, the CCTF1 strain and the aCcrK-overexpressing strain. The number of rings and total cell count :Cont:39/495; CCTF1 OE:272/592; aCcrK OE:6/1052. Scale bars: 5 μm. Statistical significance was calculated by a One-Sample t-test.*** = p value < 0.001.

## Discussion

The orderly progression of the eukaryotic cell cycle relies on precise, temporally controlled degradation of key regulatory proteins, namely cyclins, via the ubiquitin-proteasome pathway (37). Remarkably, despite the absence of canonical cyclins and CDKs, archaea of the order Sulfolobales exhibit a highly ordered cell cycle that similarly depends on proteasome-mediated proteolysis (6). Critical to this process is the AAA+ ATPase PAN, the archaeal homolog of the eukaryotic 19S regulatory particle, which unfolds proteins to facilitate their degradation by the 20S core particle (22,38,39). While it was established that the activity of PAN is crucial for the cell-cycle-dependent elimination of substrates, such as the ESCRT-III homolog CdvB, the mechanism driving the cyclic fluctuation of PAN itself remained elusive until a novel transcription factor, CCTF1, has been discovered (11). Our study confirms the key role of CCTF1 in regulation of the cell division and provides further insights into the underlying mechanism. By contrast, our data does not support the previous conclusion that aCcrK regulates the proteasome activity, leading to CdvB degradation. CCTF1-mediated transcriptional repression of *pan* occurs through specific binding to a specific palindromic sequence within the P*_pan_*. In the absence of CdvB degradation, cytokinesis dies not take place.

Our results indicate that the mechanism by which aCcrK regulates the cell division differs from that of CCTF1, contrary to the previous assumption. The overexpression of aCcrK does not appear to inhibit the proteasome-mediated degradation of CdvB, but instead seems to impede the CdvB ring formation.

Based on the results of this study and our previous findings (8,17), we propose a refined model of cell cycle regulation in Sulfolobales (Figure 8). The timely expression of cell division proteins and transcriptional repression of PAN ensures accumulation of CdvA and CdvB, which leads to the formation of the CdvB ring. Following the formation of the cytokinetic rings, further expression of the cell division proteins is repressed by aCcr1 to preclude premature reinitiation of the division ring formation. Concomitantly, repression of CCTF1 leads to the expression of *pan* and subsequent degradation of CdvB, leading to membrane constriction. Following cytokinesis, timely expression of aCcr3 ensures proper genome replication during the S phase.

**Figure 8.**
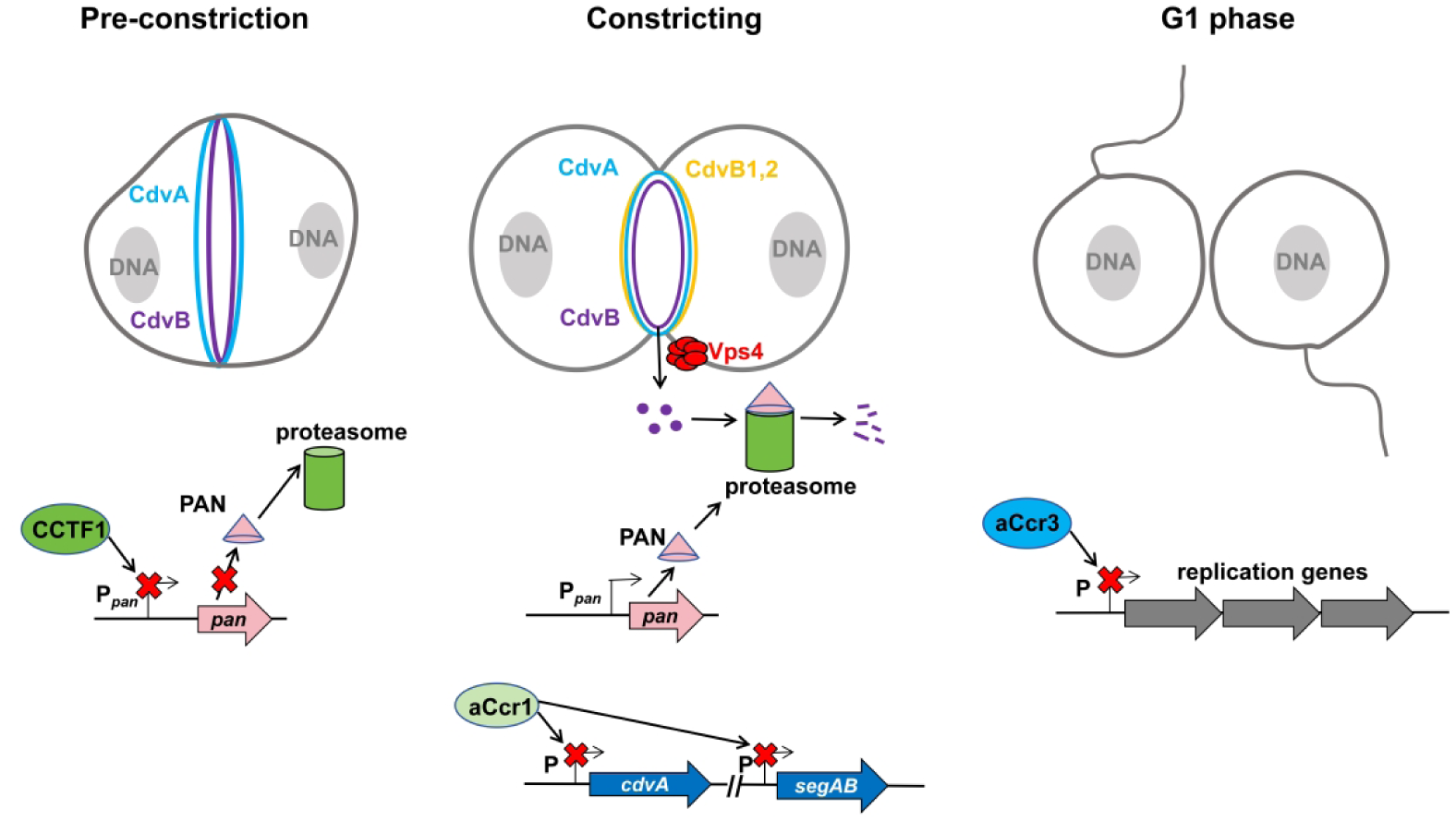
A model diagram illustrating how CCTF1 regulates cell division in Sulfolobales. Before the formation of the CdvB ring, CCTF1 suppresses the expression of PAN, thereby ensuring the assembly of the CdvB ring. During the contraction phase of the cell division ring, the decrease in CCTF1 expression lifts the inhibition of PAN, thereby initiating the proteasomal degradation of cell division proteins. Meanwhile, the expression of aCcr1 ensures that cell division and chromosome segregation do not restart. In newly born G1-phase cells, the expression of aCcr3 ensures that replication-related genes are activated only once.

The C-terminal region of CdvB, specifically the broken-winged helix domain, mediates the binding to CdvA on the one hand and functions as a potent degradation signal, on the other hand (11). This finding suggests that the interaction by which CdvB is recruited to the division plane also primes it for destruction (11). However, the acceleration of CdvB degradation during cytokinesis is not solely dependent on this intrinsic signal. It requires the concurrent de-repression of *pan*, as cells enter into the division phase. By contrast, aCcrK does not appear to function in the regulation of the proteosome-dependent CdvB degradation.

The CCTF1-PAN paradigm reveals a striking case of convergent evolution in cell cycle regulation: although archaea and eukaryotes employ entirely different molecular players for the regulation of cell cycle progression, the outcome is similar. In eukaryotes, the timing of proteolysis is primarily determined by post-translational modifications that confer recognition specificity (such as APC/C-mediated ubiquitination)(37). In contrast, Sulfolobales appear to have evolved a simpler transcription-based regulatory mechanism, in which transcription of the proteasome activating factor serves as the primary oscillating variable. This mechanism could represent the ancestral cell cycle regulation via protein degradation, which predates the evolution of complex ubiquitin ligase networks in more complex organisms. Potentially, the ancestral role of the proteasome activity in cell division could have been regulated at the transcriptional level, whereas the layered complexity of ubiquitination emerged during later stages of evolution, enabling more precise and substrate-specific regulation.

### Data avilability

All data supporting the findings of this study are available within the article and its Supplementary Information, or from the corresponding author upon reasonable request.

## Acknowledgements

We would like to thank members of the CRISPR and Archaea Biology Research Center for helpful discussions and technicians from the Core Facilities for Life and Environmental Sciences, State Key Laboratory of Microbial Technology of Shandong University for assistance. This work was supported by National Natural Science Foundation of China [32370033 and 32393973] to Y.S., Postdoctoral fellowship Program of CPSF (GZC20231471) and Postdoctoral Innovation Project of Shandong Province (SDCX-ZG-202400122) to Y.Y., and the State Key Laboratory of Microbial Technology Open Projects Fund [Project NO. M2023-20, M2025-12] to Y.S. We thank Jing Zhu, Zhifeng Li, Guannan Lin, and Jingyao Qu of the Core Facilities for Life and Environmental Sciences, State Key Laboratory of Microbial Technology of Shandong University for their helps in protein analysis, flow cytometry and microscopy.

## Author contributions

**Yunfeng Yang**: Conceptualization; Data curation; Formal analysis; Investigation; Visualization; Methodology; Supervision; Funding acquisition; Project administration; Writing-original draft and editing. **Zixin Geng**: Investigation; Data curation; Methodology.**Shikuan Liang**: Investigation; Methodology. **Haodun Li**: Investigation; Methodology.**Fan Zhou**:Investigation; Methodology. **Junfeng Liu**: Methodology. **Mart Krupovic**: Formal analysis; Writing-review and editing. **Yulong Shen**: Supervision; Funding acquisition; Project administration; Writing-review and editing.

## SUPPLEMENTARY FIGURES AND TABLES

**Figure S1.**
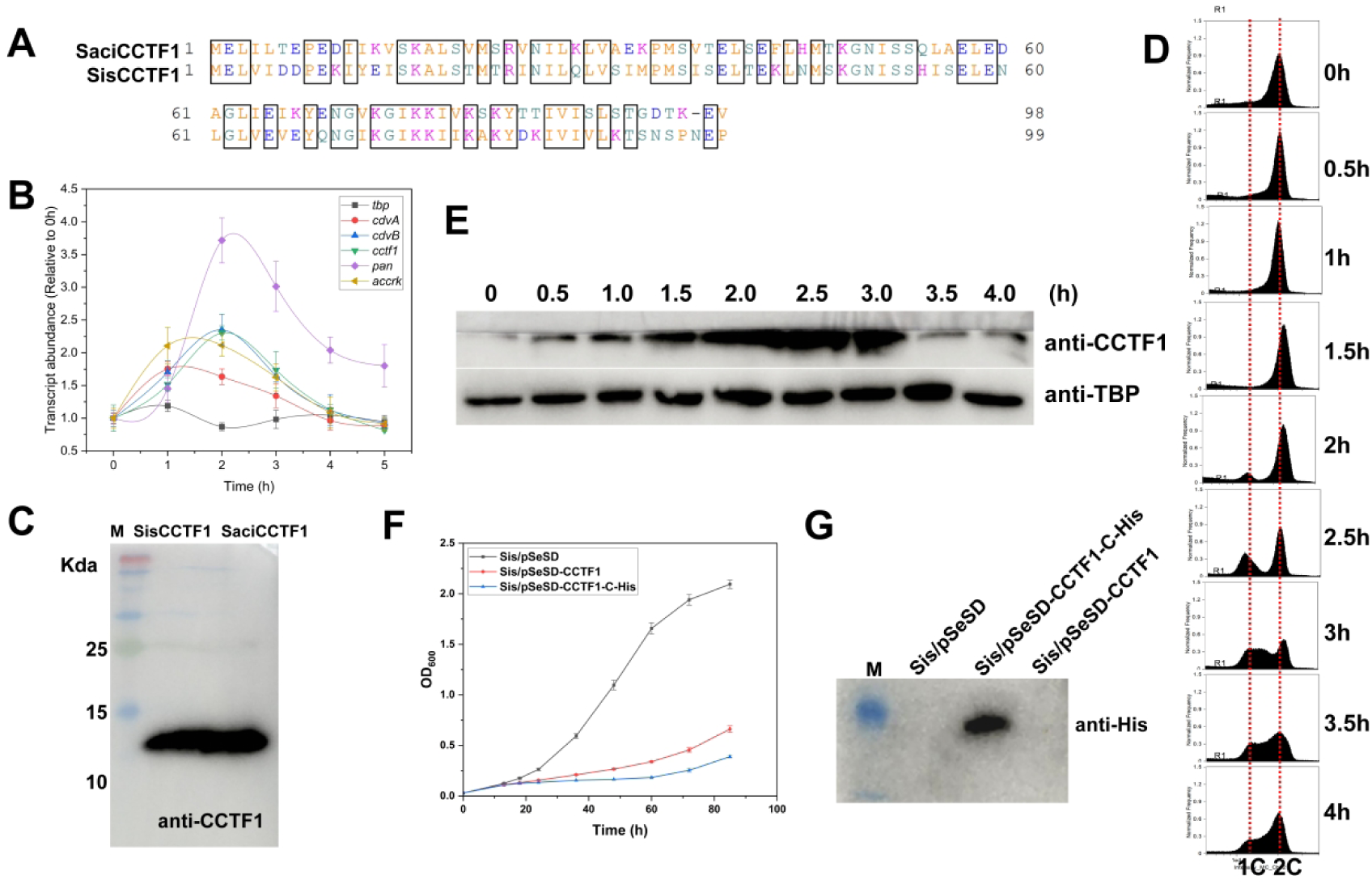
(**A**) Sequence alignment of SaciCCTF1(Saci_0800) and its homolog SisCCTF1(SiRe_1806) in *Sa. islandicus*. (**B**) Expression patterns of *siscctf1* and related genes (*cdvA*, *cdvB*, *pan*, *aCcrK*). *tbp* was used as a control. The profiles were obtained based on transcriptomic data of synchronized cells (8). (**C**) The antibody targeting SisCCTF1 *in vitro* exhibits equivalent efficacy towards the CCTF1 proteins. The amount of 1.0 ng SisCCTF1 or SaciCCTF1 recombinant protein was loaded for the analysis. (**D**) Flow cytometry profiles of samples of a synchronized *S. acidocaldarius* DSM639 culture. The cells were synchronized at G2 phase after treated with 6 mM acetic acid for 4 h before released by removing the acetic acid. The cultures were collected at different time points (0, 0.5,1.0, 1.5, 2.0, 2.5, 3.0, 3.5 and 4.0h) and subjected to flow cytometry analysis. “1C” and “2C” represent one and two chromosomes, respectively. (**E**) Western blotting verification of the expression of CCTF during different cell cycle phases of the *S. acidocaldarius* DSM639. TBP was used as a loading control. (**F**) Cells expressing the C-terminal His-tagged SaciCCTF1 display the same phenotype as that expression the untagged protein. (**G**) Validation of CCTF1 expression by Western blotting using anti His antibody.

**Figure S2.**
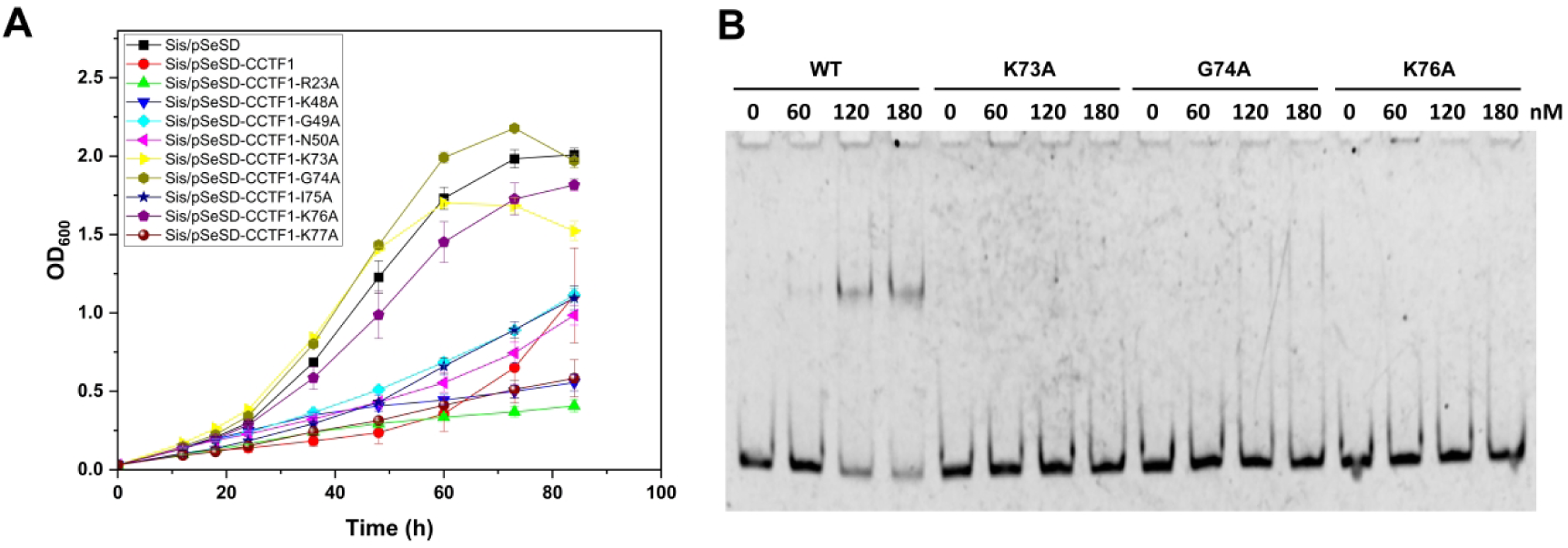
The residues K73, G74 and K76, located in the β-hairpin, are crucial for the DNA-binding ability of *Sis*CCTF1. (**A**) Growth curves of the strains overexpresing CCTF1 and alanine mutants of conserved residues. (**B**) EMSA for the binding capacity of the K73A, G74A and K76A mutants. P*_pan_* was used as the substrates.

**Figure S3.**
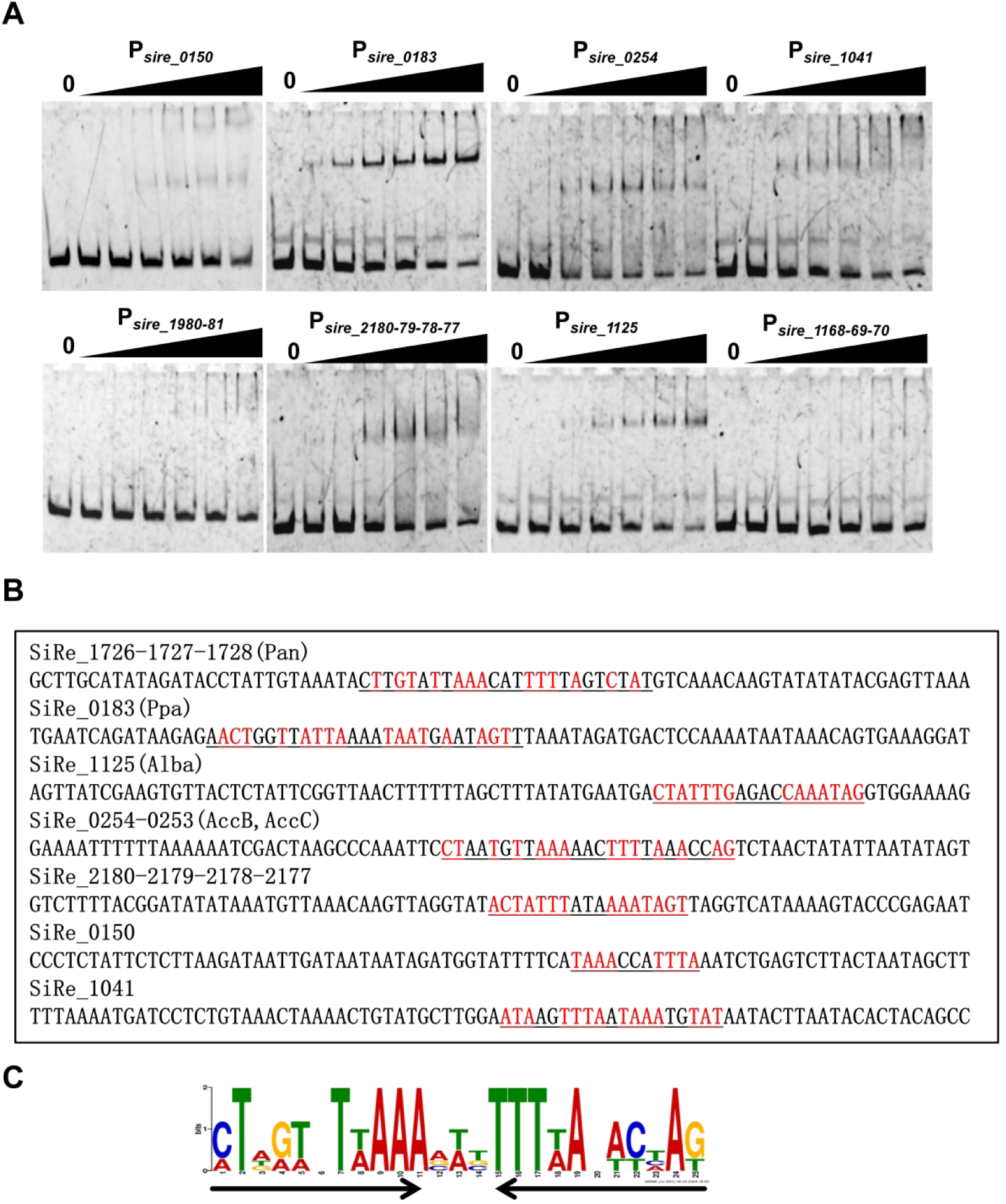
(**A**) EMSA analysis of the binding ability of CCTF1 of the promoters of highly downregulated genes in the overexpression sstrain. The amount of promoter used was 10 ng. The protein concentrations were increased in the following order: 0, 30, 60, 120, 180, 240, and 300 nM. (**B**) Sequence feature of the promoters which CCTF1 can bind in vitro.(**C**)A conserved motif identified in the promoter of the highly repressed genes that CCTF1 can bind their promoters in vitro by MEME server with the default setting. The height of the letter indicates the relative similarity to that of consensus one.

**Figure S4.**
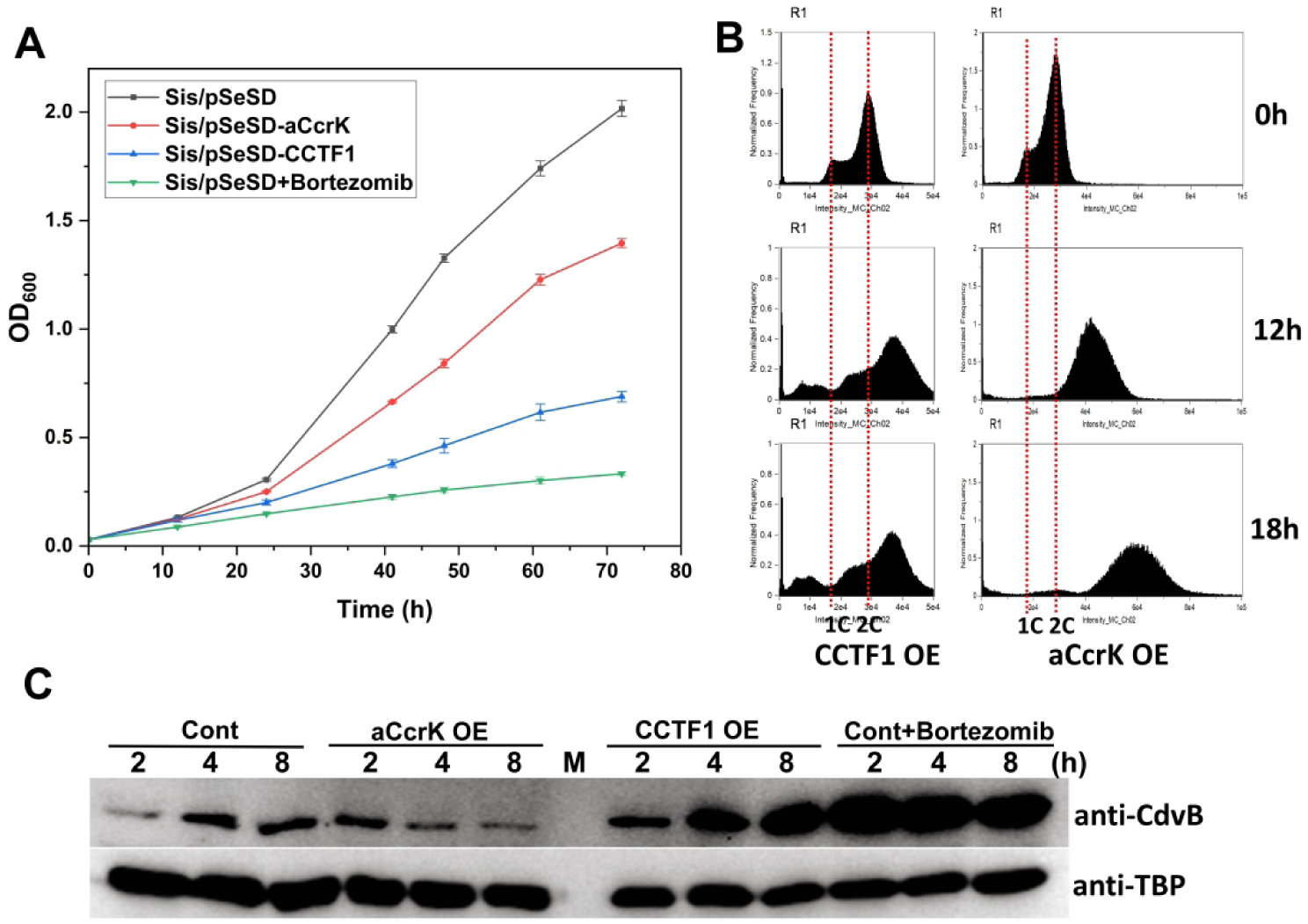
(**A**) Comparison of growth curves between aCcrK-overexpressing, CCTF1-overexpressing, and wild-type strains treated with a proteasome inhibitor (final concentration 1 μM Bortezomib). (**B**) The CCTF1- and aCcrK- overexpressing strains display completely different flow cytometric profiles as the duration of arabinose induction increased.(**C**) Changes in CdvB protein levels at different time points in control, aCcrK-overexpressing, CCTF1-overexpressing, and proteasome inhibitor-treated strains. TBP was used as an internal control.

**Figure S5.**
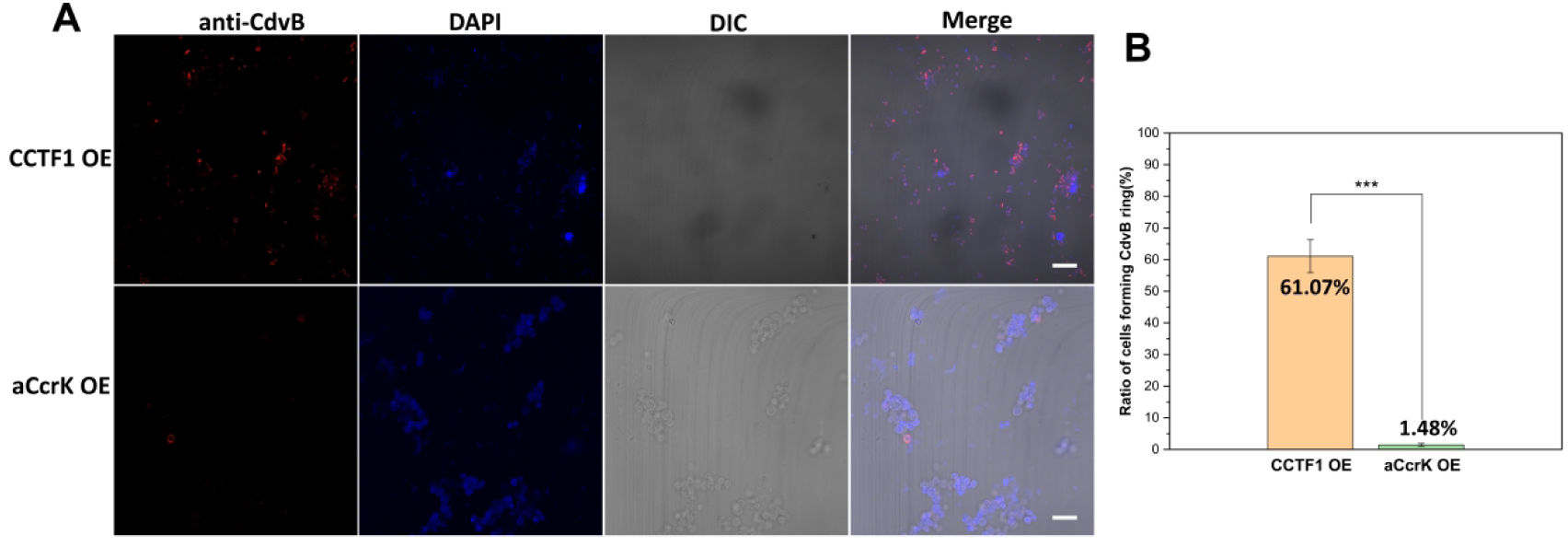
(**A**) At 20 hours post arabinose induction, there was a marked more CdvB rings were formed in the CCTF1-overexpressing cells than the aCcrK-overexpressing cell. Scale bars: 5 μm. (**B**) Statistical analysis of the proportion of CdvB ring formation. The number of rings and total cell counts : CCTF1 OE:210/347; aCcrK OE:4/259. Scale bars: 5 μm. Statistical significance was calculated by a One-Sample t-test.*** = p value < 0.001.

**Table S1.**
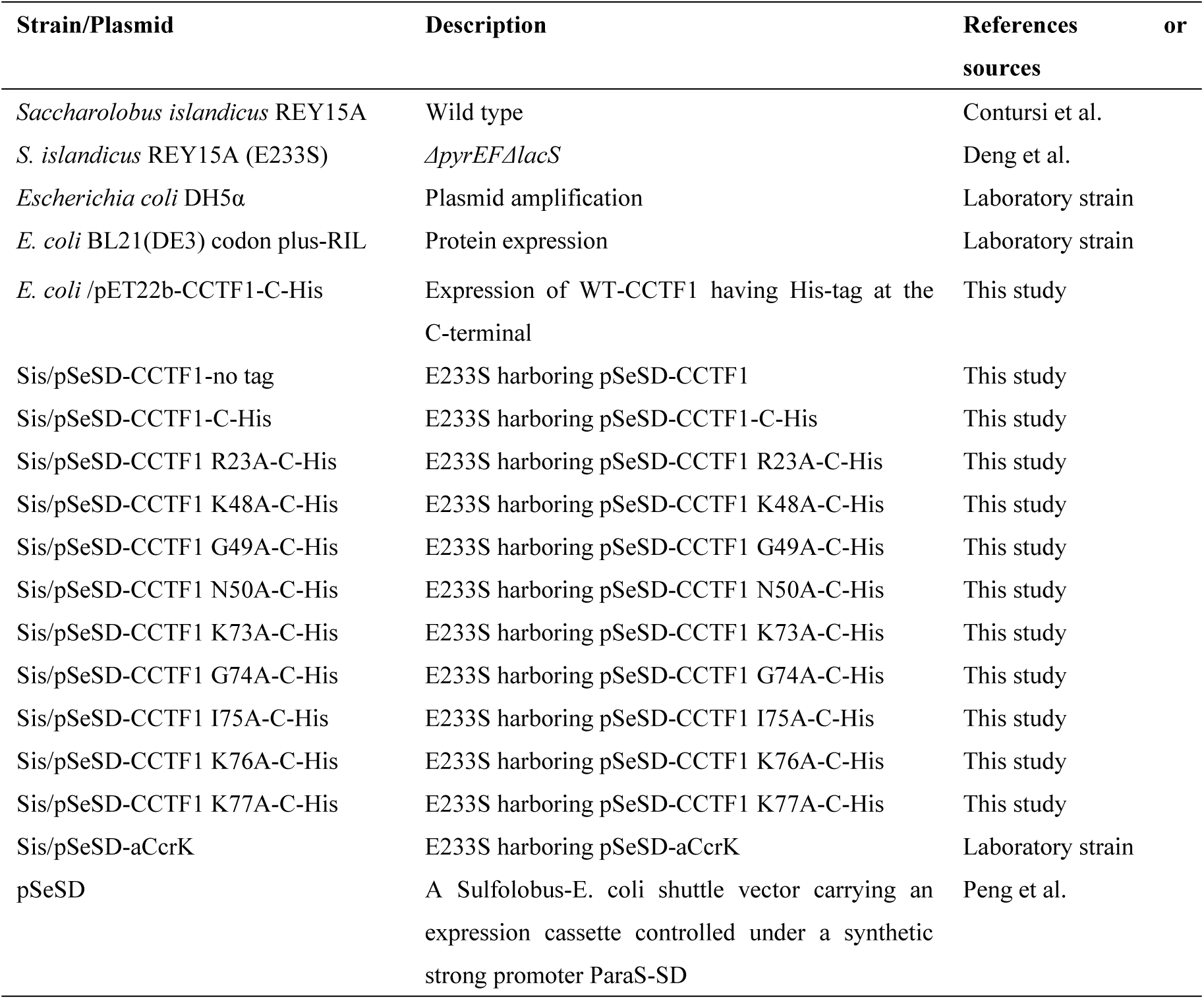
Plasmids and strains used in this study.

**Table S2.**
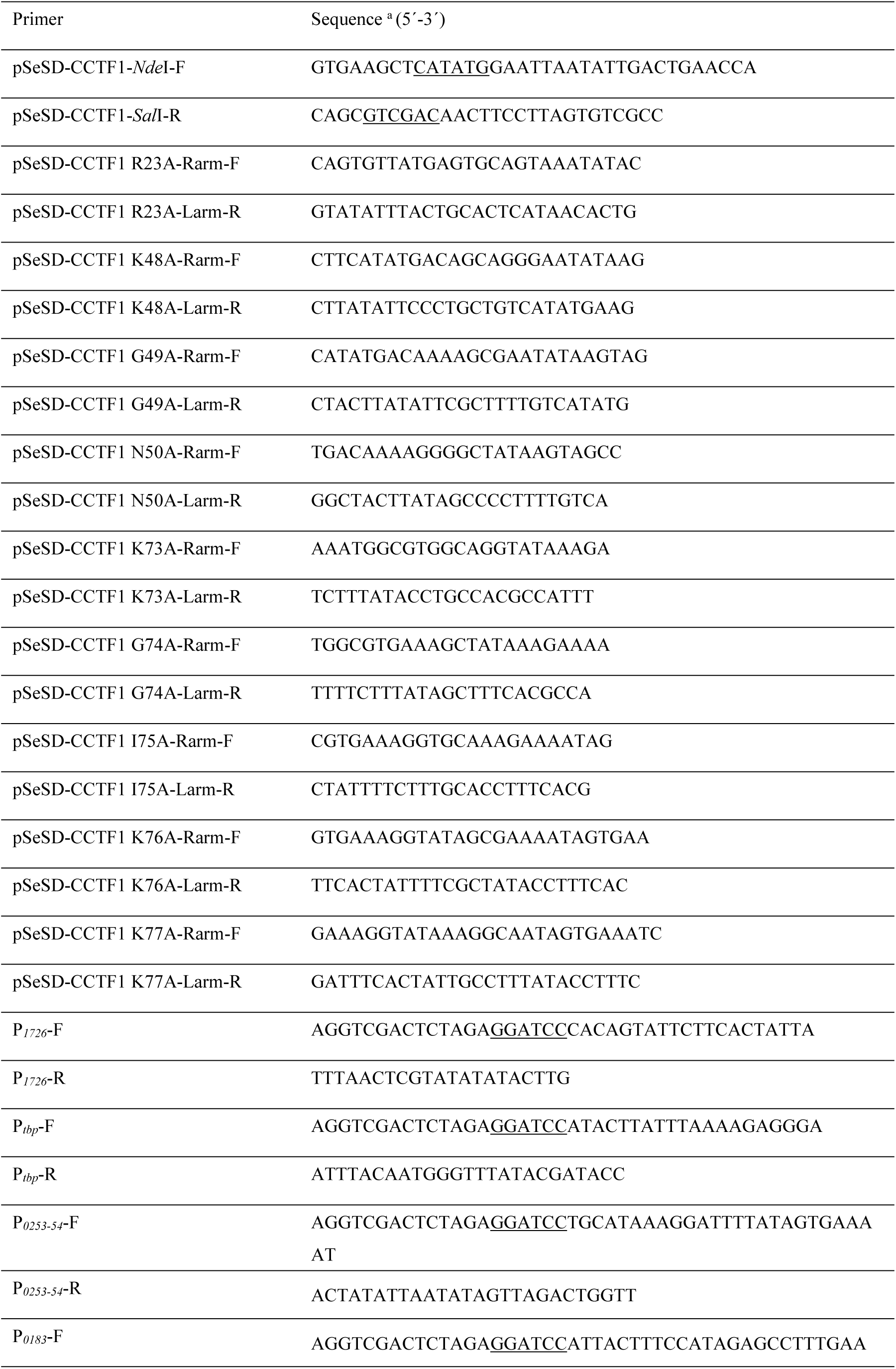
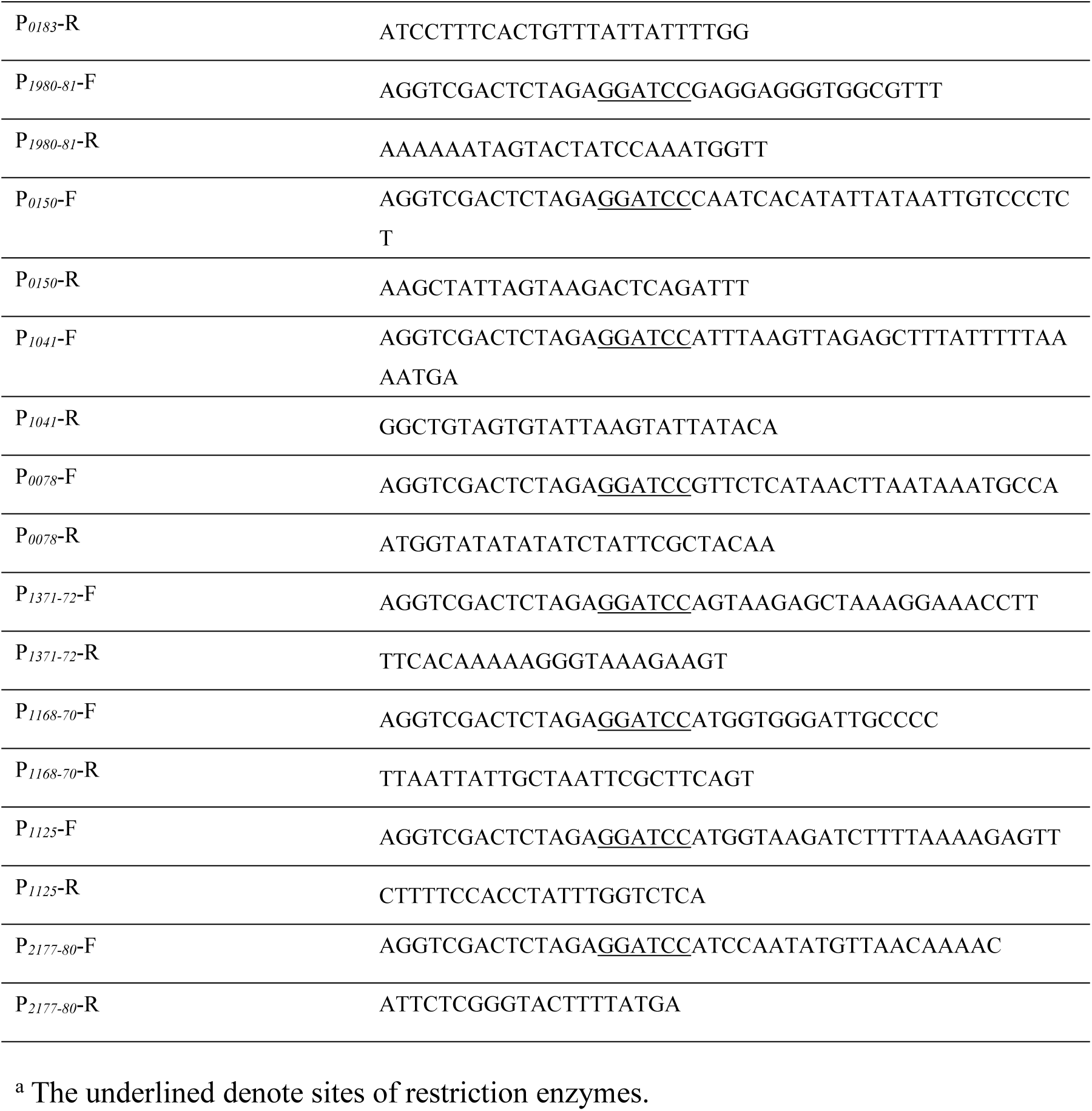
Oligonucleotides used as primers in this study.

